# Dopaminergic signaling regulates microglial surveillance and adolescent plasticity in the frontal cortex

**DOI:** 10.1101/2024.03.08.584167

**Authors:** Rianne Stowell, Kuan Hong Wang

## Abstract

Adolescence is a sensitive period for frontal cortical development and cognitive maturation. The dopaminergic (DA) mesofrontal circuit is particularly malleable in response to changes in adolescent experience and DA activity. However, the cellular mechanisms engaged in this plasticity remain unexplored. Here, we report that microglia, the innate immune cells of the brain, are uniquely sensitive to adolescent mesofrontal DA signaling. Longitudinal *in vivo* two-photon imaging in mice shows that frontal cortical microglia respond dynamically to plasticity-inducing behavioral or optogenetic DA axon stimulation with increased parenchymal and DA bouton surveillance. Microglial-axon contact precedes new bouton formation, and both D1 and D2-type DA receptors regulate microglial-bouton interactions and axonal plasticity. Moreover, D2 antagonism in adults reinstates adolescent plasticity, including increased microglial surveillance and new DA bouton formation. Our results reveal that DA signaling regulates microglial surveillance and axonal plasticity uniquely in the adolescent frontal cortex, presenting potential interventions for restoring plasticity in the adult brain.

## Introduction

Adolescence is increasingly viewed as a sensitive period for the structural and functional development of the frontal cortex^1,2^. Psychiatric disorders, such as schizophrenia and attention deficit hyperactivity disorder (ADHD), frequently manifest in adolescence^3^. This coincides with the maturation of the dopaminergic (DA) mesofrontal circuit, which connects the midbrain motivation center to the frontal cortical cognitive control center^4–6^. Importantly, disruptions in the development and function of the frontal cortical dopamine system are key characteristics of neurodevelopmental psychiatric disorders^7,8^. Despite our long-standing recognition of the disease relevance of frontal DA dysfunction, little is known about the cellular mechanisms involved in the adolescent plasticity of frontal DA circuits.

The adolescent mesofrontal circuit is malleable in response to changes in behavioral experience and DA activity^9,10^. Frontal DA axons exhibit robust outgrowth of boutons, the main release sites for DA^6,11^, in response to rewarding behavioral experience such as wheel-running in mice^9^. Furthermore, directly increasing the phasic activity of ventral tegmental area (VTA) DA neurons, a pattern of activity typically associated with reward^12^, induces frontal DA bouton formation, as well as functional changes in frontal cortical activity and psychomotor behavior^9^. Adolescent stimulation of VTA DA neurons or frontal DA projections to drive mesofrontal plasticity rescues frontal circuit and behavioral deficits in multiple genetic mouse models of neurodevelopmental disruption^13^. However, mesofrontal DA circuit plasticity is unique to adolescence, as phasic VTA stimulation alone cannot elicit these effects in adult animals^9^, suggesting that the adolescent frontal cortex is uniquely permissive of activity-dependent changes. Interestingly, adult plasticity can be reinstated by pairing VTA stimulation with the pharmacological blockade of the D2-type DA receptor (D2R)^9^. However, it remains unknown what cellular players are engaged in the mesofrontal plasticity during adolescence and the reinstating of this plasticity in adulthood.

In pursuit of candidate regulators of mesofrontal DA plasticity, we investigated the adolescent microglial response to DA signaling. Beyond their roles in pathological conditions, microglia, the innate immune cells of the central nervous system (CNS), can modulate neural circuit development and activity-dependent plasticity^14–18^. Recently, microglia are also being recognized as a heterogeneous population of cells with regional specificity in both gene expression and responsiveness to CNS signals^19^. The differences between subcortical and cortical microglia have recently been examined through RNA sequencing studies, but the developmental and spatial diversity of microglia across cortical regions remains unexplored^20^. Importantly, while microglia respond to a few neurotransmitters such as norepinephrine^15,21^, GABA^22^ and glutamate^23^ in several brain regions, it is unknown if microglia respond to DA signals *in vivo.* Previous *in vitro* work suggests that microglia express DA receptors and can respond to DA signaling^24^. Recent studies in the basal ganglia found that microglia exhibit region-specific phenotypes^25^ and influence neuronal DA receptor expression in the adolescent nucleus accumbens^18^. However, critically, previous research has not investigated whether microglia respond to DA *in vivo* or interact with the frontal DA circuit. Thus, evaluating microglia as a potential cellular player in frontal DA circuit plasticity may provide key insight into plasticity mechanisms and inform future therapeutic strategies.

Here, we investigated if frontal cortical microglia respond to DA signals from the mesofrontal circuit. We show that *in vivo* adolescent microglia respond to the activity of the mesofrontal circuit elicited by both wheel running and direct phasic activation of the mesofrontal DA axons. Our findings depict a biphasic microglial response to phasic mesofrontal activation, characterized by an initial retraction of microglial processes during stimulation and a subsequent sustained increase in parenchymal surveillance, which includes enhanced contact with axonal elements. Critically, during axonal surveillance, we find that microglial contacts along the axon backbone preceded the formation of new boutons. Furthermore, we demonstrate that perturbing DA signaling through pharmacological manipulation of either D1 or D2 receptors can eliminate the biphasic response of frontal cortical microglia to phasic DA activity and block new bouton formation. Finally, we show that reinstating plasticity in adult mice by D2 antagonism results in an adolescent-like microglial DA response and increased surveillance of DA boutons. Our work shows that adolescent frontal cortical microglia are exquisitely sensitive to mesofrontal DA signaling and that their dynamics are tightly coupled to both phasic mesofrontal activity and subsequent DA bouton outgrowth in the frontal cortex.

## Results

### Adolescent wheel running promotes DA bouton formation and microglial process outgrowth in the frontal cortex

In order to determine if microglia respond to wheel-running induced mesofrontal plasticity, we utilized Cx3cr1^GFP^ mice crossed to a Th-Cre line so that microglia would be labeled with GFP and DA projections from the VTA could be specifically labeled through AAV-CAG-FLEX-tdTomato injected into the VTA^9,13,15^ (Fig. 1a). We used Th-Cre rather than DAT-Cre as mesofrontal dopamine neurons have much stronger TH expression but little DAT expression^26,27^, which limits the utility of the DAT-Cre line for our target dopaminergic neuronal population. To verify Th-Cre was the appropriate choice we confirmed that our virus transduction was limited to the VTA injection site (Supplementary Figure 1a,b). We also evaluated DAT-Cre/Ai14 mice and found that there was a lack of DA axon labeling in the frontal cortex as well as spurious ectopic neuronal labeling^28^ (Supplementary Figure 1c,d), suggesting that these mice would not be efficacious for our experimental purposes.

**Figure 1.**
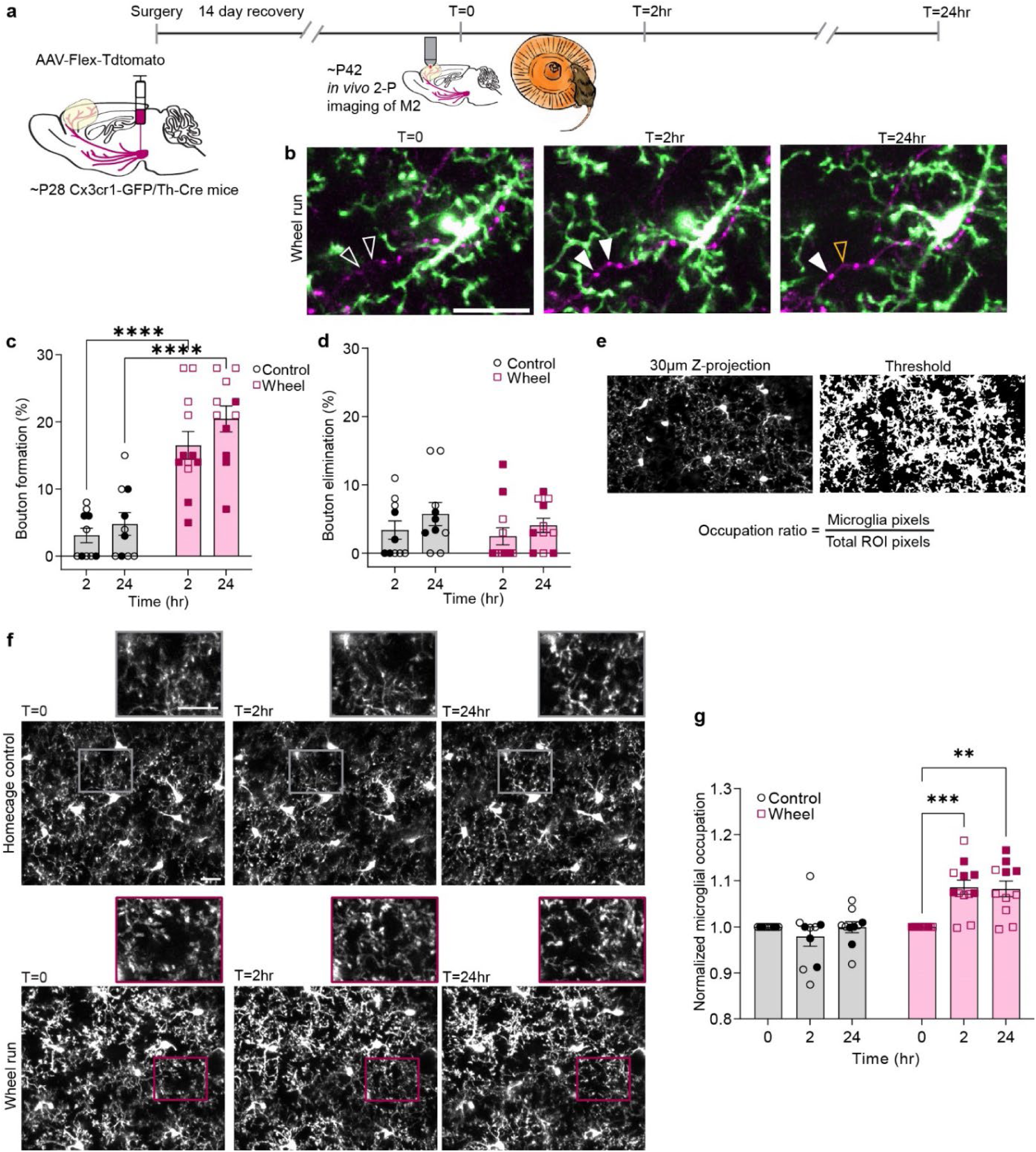
Adolescent wheel running promotes DA bouton formation and microglial process outgrowth. (**a**) Timeline of surgical and imaging procedures. (**b**) Example of microglia (green) and axons (magenta) at t=0 (pre-run), after 2hr of wheel running, and 24hr later. Hollow arrowheads in t=0 denote areas where new boutons form post-wheel run (solid arrowheads), with the hollow orange arrowhead denoting a lost bouton at t=24hr. (**c**) 2hrs of wheel running promotes formation of new boutons (n: control=10, wheel=12 mice Mixed-effects model, Fixed effects [type III], Run status p<0.0001, F (1,20)=36.84, Šídák’s multiple comparisons Control v. Wheel, 2hr p<0.0001 & 24hr p<0.0001) (**d**) Wheel running does not affect the rate of bouton elimination (n: control=10, wheel=12 mice, Mixed-effects model, Fixed effects [type III], Run status p=0.5080, F(1,20)=0.5453, Šídák’s multiple comparisons test Control v. Wheel, 2hr p=0.8682 & 24hr p=0.7674) (**e**) Visual representation of microglial occupation analysis. (**f**) Example 30µm z-projections of microglia. Rectangle pop-outs denote representative areas of microglial process occupation. (**g**) Wheel running increases microglial occupation of the parenchyma. (n: control=10, wheel=12 mice, Mixed-effects model, Fixed effects [type III], Time x Run status p<0.0001, F(2,39)=12.17, Tukey’s multiple comparisons wheel 0 v. 2hr p= 0.0004, 0 v. 24hr p= 0.0017). Scale bars 20µm. Graphs show mean ± S.E.M **p<0.01, ***p<0.005, ****p<0.0001. Individual points represent individual animals with females as hollow symbols and males as solid symbols.

For all experiments, we used a chronic cranial window approach to allow for repeated imaging and to avoid the potential confound of the effects of anesthesia on microglial dynamics^15^. We selected the M2 area of the frontal cortex for imaging as it is optically accessible without the need for an invasive approach, which would disrupt microglia dynamics. In addition, this area receives robust dopaminergic innervation for *in vivo* imaging^4,9,29^. Our mice were imaged in the middle of adolescence between P35-P49^30^ when the mesofrontal circuit is still undergoing maturation^4^ and remains plastic^9^. We also confirmed that chronic cranial window preparations did not impact wheel running behavior (Supplemental figure 2a).

We found that after two hours of wheel running, both male and female mice showed increased rates of bouton formation and that this increase in boutons was maintained at 24hrs (Fig. 1b,c. Mixed-effects model, Fixed effects [type III], Run status p<0.0001). The increased bouton formation was not accompanied by a change in elimination (Fig. 1d Mixed-effects model, Fixed effects [type III], Run status, p=0.4069), demonstrating that in both male and female adolescent mice wheel running generates a net gain in DA boutons. Interestingly, although both male and female mice had increased bouton formation post wheel running, further analysis found that female mice had significantly higher bouton formation rates than males (Mixed-effects model, Fixed effects [type III], Sex x Run status p=0.0221). We did not find a significant difference between male and female run distance (Supplementary Figure 2b), nor a significant correlation between run distance and bouton formation rate (Supplementary Figure 2c).

On the same timescale as the bouton changes, we found significant changes in microglial arborization. We used occupation, the ratio of microglial occupied pixels out of the total pixels in the field of view, as a measure of the degree of microglial arborization. Microglia with more elaborate arbors tend to occupy more of the parenchyma^15^ (Fig.1e). We found that microglial occupation of the parenchyma significantly increased after 2hrs of wheel running and that microglia remained in this more ramified state at 24hrs (Fig. 1f,g. Mixed-effects model, Fixed effects [type III], Time x Run status p<0.0001). Thus, 2hrs of wheel running, which is known to drive phasic VTA activity^31^, is sufficient to generate both DA bouton formation and concomitant increases in microglial arborization.

### Wheel running does not drive microglial process outgrowth in the adult frontal cortex or in the adolescent visual cortex

Our previous work demonstrated that mesofrontal plasticity from wheel running was unique to adolescence, and that adult mice did not show a significant increase in DA bouton formation after 2hrs of wheel running^9^. Interestingly, our adult mice ran significantly more in the 2hrs than the adolescent mice (Fig 2.a, unpaired t-test p<0.005), but neither male nor female adult mice showed any changes in bouton formation or elimination after wheel running (Fig. 2b,c. Mixed-effects model, Fixed effects [type III], Run status, formation p=0.1443 and elimination p=0.6900). Critically, the adult mice also showed no changes in microglial arborization (Fig. 2d,e Mixed-effects model, Fixed effects [type III], Run status, p=0.4076) suggesting that the effect of wheel-running on microglia in adolescence was not simply a product of exercise, particularly because the adults actually ran further than the adolescents.

**Figure 2.**
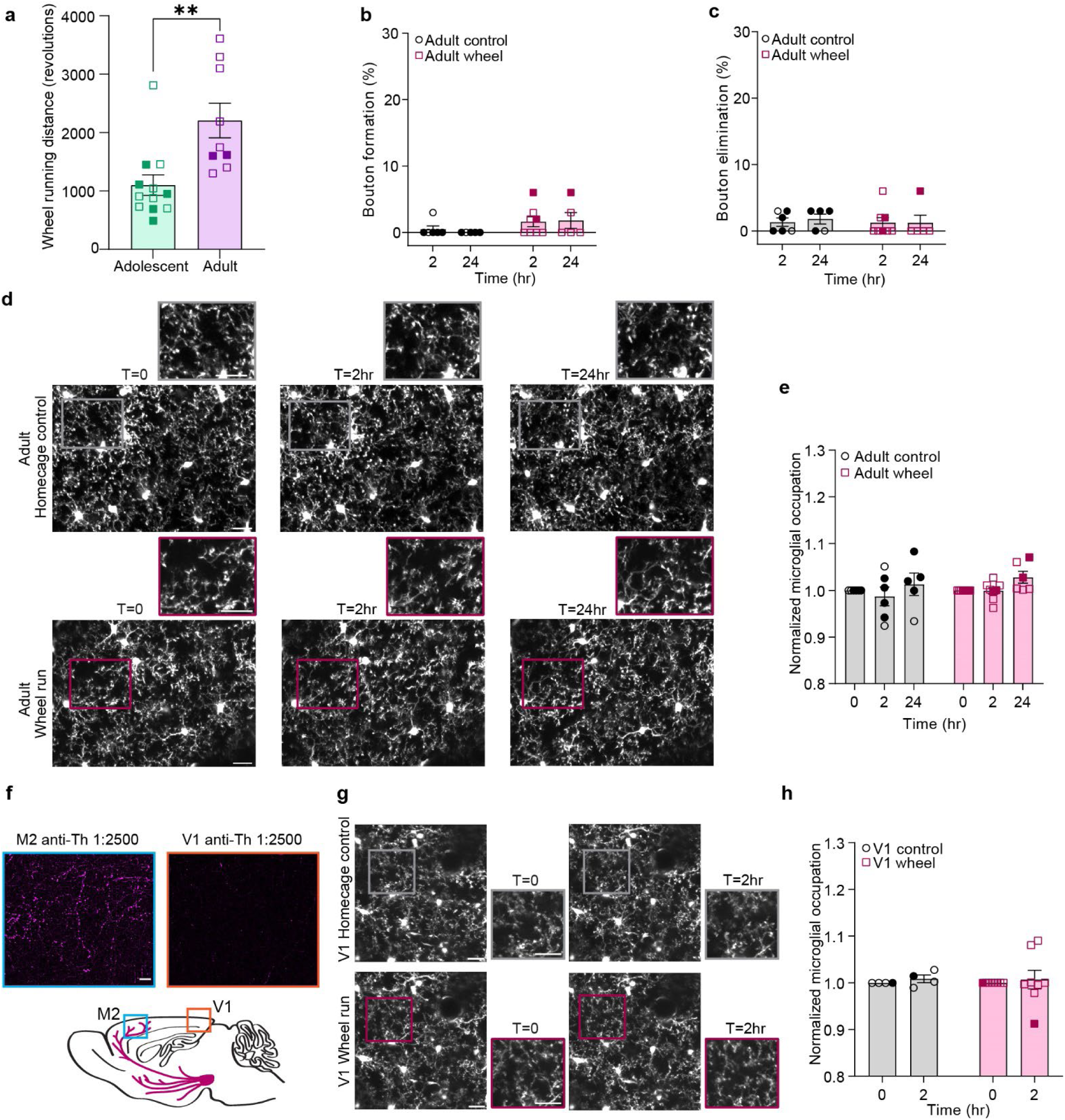
DA bouton outgrowth and the concomitant microglial process outgrowth are unique to the adolescent frontal cortex. (**a**) Adult mice run more than adolescent mice during 2hrs on a running wheel (n: adolescent=12, adult=9 mice, two-sided unpaired t-test p=0.0032, t(19)=3.375).(**b**) Wheel running does not drive DA bouton outgrowth in adult mice (n: control=6, wheel=8 mice, Mixed-effects model, Fixed effects [type III], Run status p=0.1443, F(1,12)=2.440, Šídák’s multiple comparisons Control v. Wheel 2hr p=0.25 & 24hr p=0.25).(**c**) Wheel running does not drive DA bouton elimination in adult mice (n: control=6, wheel=8 mice, Mixed-effects model, Fixed effects [type III], Run status p=0.69, F(1,20)=0.1638, Šídák’s multiple comparisons Control v. Wheel 2hr p=0.9964 & 24hr p=0.8742).(**d**) Example 30µm z-projections of adult microglia in the M2 frontal cortex. Rectangle pop-outs denote representative areas of microglial process occupation. (**e**) Wheel running does not increase microglial occupation in adult mice (n: control=6, wheel=8 mice, Mixed-effects model, Fixed effects [type III], Run status p=0.4076, F(1,12)=0.7364, Šídák’s multiple comparisons Wheel 0 v. 2hr p=0.9786 & 0 v. 24hr p=0.1115) (**f**) Anti-tyrosine hydroxylase staining of M2 and V1 at a 1:2500 dilution, which captured dopaminergic but not noradrenergic fibers. Below the staining is a diagram highlighting the substantial dopaminergic innervation of M2 compared to V1, which receives few dopaminergic projections. (**g**) Example 30µm z-projections of microglia. Rectangle pop-outs denote representative areas of microglial process occupation. (**h**) Wheel running does not increase microglial occupation in V1 (n: control=4, wheel=6 mice, Two-way repeated measures ANOVA, p=0.9538, F(1,10)=0.003523). Scale bars 20µm. Graphs show mean ± S.E.M **p<0.01. Individual points represent individual animals with females as hollow symbols and males as solid symbols.

To further test the specificity of wheel-running induced adolescent microglia changes to the frontal cortex, we evaluated mice with windows over the primary visual cortex (V1), which lacks significant DA innervation as compared to M2^32^(Fig. 2f). We found that in V1 microglia did not respond to wheel running (Fig. 2g,h Two-way repeated measures ANOVA, p=0.9538). Thus, the effect of wheel-running on microglia is specific to adolescence and the cortical region receiving major DA input from the VTA.

### Microglial surveillance decreases during DA axon stimulation and then increases post-stimulation in adolescent frontal cortex

After observing the parallel changes in DA boutons and microglia in the frontal cortex induced by wheel running, we wanted to further determine if microglia respond to direct changes in DA neuron activity. To this end, we injected AAV2/9-CAG-Flex-ChR2-tdTomato into the VTA of Th-Cre mice to allow for optogenetic stimulation of DA pathways. With direct light stimulation of mesofrontal DA axons over the cranial window, we confirmed the efficacy of this stimulation method by conducting two-photon calcium imaging of the cortical neural response to phasic DA stimulation^13,33^ (Supplemental Fig. 3).

Based on the tight temporal control of stimuli enabled by our optogenetic approach, we characterized the time course of the microglial response to phasic DA axon stimulation (473nm, 20mW output, 50Hz pulse train, 3ms/pulse, 10 pulses/train, 1 train/min for 10min) by imaging microglia every minute between our pulses of light stimulation (Fig.3a). With these sequential images, we assayed microglial surveillance, which captures microglial occupation accumulated over time. By collapsing images across time, we can quantify all the areas microglia surveyed within our sequence of images^15^ (Fig. 3a). We found that in response to DA axon activation, microglia reduced their surveillance of the surrounding parenchyma as compared to their baseline surveillance without stimulation (Fig. 3b,c paired t-test, p<0.01, Supplemental Videos 1 and 2). Upon cessation of stimulation (from 10 to 90min after the start of the first stimulation), we found that microglial surveillance rapidly recovered and significantly surpassed the level of surveillance in unstimulated animals (Fig. 3d,e. paired t-test, p<0.001, Supplemental Videos 3 and 4).

**Figure 3.**
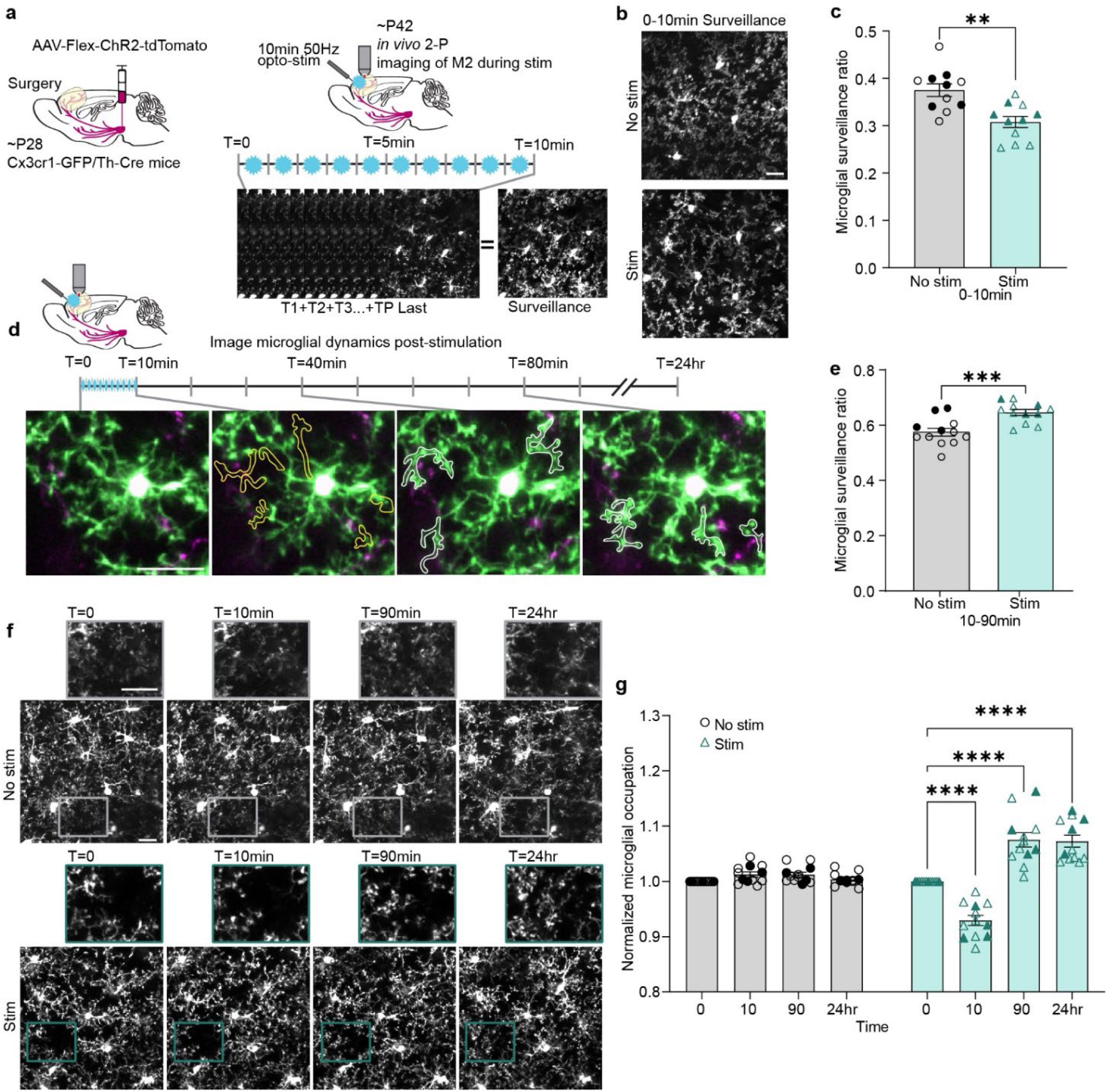
Microglia have a biphasic response to optogenetic stimulation of DA axons. (**a**) Diagram of stimulation and image collection parameters. (**b**) Example microglial surveillance with and without opto-genetic stimulation. (**c**) Microglial surveillance is reduced during phasic stimulation of the mesofrontal DA axons (n=11 mice, two-tailed paired t-test, p=0.0015, t(10)=4.324). (**d**) Example of individual microglial process retraction and extension during imaging. Immediately after stimulation, microglial processes occupy less parenchyma (retracted processes outlined in yellow). Over the course of 90min, microglia extend processes (white outline marking process outgrowth) toward axons (magenta). (**e**) After optogenetic stimulation, microglia increase surveillance of the parenchyma (n=12 mice, two-tailed paired t-test, p=0.0003, t(11)=5.213). (**f**) Representative images of microglial occupation changes pre-stim (T=0), immediately after stimulation (T=10min), 90min later, and 24hr after phasic stimulation, as well as paired no-stim timepoints (rectangular pop-outs highlight changes in microglial processes). (**g**) DA axon stimulation drives a biphasic microglial occupation change with an initial reduction followed by a prolonged increase in parenchyma occupation (n=12 mice, two-way repeated measures ANOVA time x stimulation p<0.0001, F(3,33)=47.02, Holm-Šídák’s multiple comparisons Stim 0 v. 10min p<0.0001, 0 v. 90min p<0.0001, & 0 v. 24hr p<0.0001). Scale bars 20µm. Graphs show mean ± S.E.M **p<0.01, ***p<0.005, ****p<0.0001. No stim and stim experiments were conducted within the same animals. Individual points represent individual animals with females as hollow symbols and males as solid symbols.

We then further assessed the changes over the entire course of stimulation and imaging by calculating microglial occupation at 0min, 10min, 90min, and 24hrs. By examining occupation at these individual time points, we observed an initial dip in occupation from 0 to 10min, corresponding with the stimulation period, followed by a significant increase in occupation by 90min which was sustained to 24hrs (Fig. 3f,g two-way repeated measures ANOVA time x stimulation p<0.0001). This was not an effect of the light exposure itself, as a cohort expressing only td-Tomato while receiving light stimulation did not show these responses (Supplemental Fig. 4a-c).

Comparing our optogenetic stimulation to our wheel running cohort, we see a parallel in the post-treatment microglial response pattern—after phasic stimulation of M2 DA axons or wheel running, microglia undergo process elaboration and maintain this increased parenchymal occupation for 24hrs post-stimulation. Critically, this increase in microglial surveillance in both wheel running and optogenetic stimulation conditions coincides with the time period when new DA boutons formed with wheel running. Thus, we next evaluated what relationship increased microglial surveillance may have to DA bouton plasticity with the optogenetic approach.

### Microglia contact the axonal sites for newly formed boutons and increase surveillance of stable boutons after adolescent DA axon stimulation

We first confirmed that, as in the wheel running paradigm, optogenetic phasic DA axon stimulation resulted in increased bouton formation at 2hrs, which was sustained out to 24hrs without a concomitant change in elimination (Fig. 4a-c; formation two-way repeated measures ANOVA stimulation p<0.0001; elimination two-way repeated measures ANOVA, stimulation, p=0.8194). To track microglial contacts with boutons, we identified stable boutons (boutons present throughout all the imaging time points), new boutons (boutons appearing along the axon backbone after t=0), and eliminated boutons (boutons present at t=0 but disappearing prior to or by 24hrs). For new or eliminated boutons, microglial contacts with the axon backbone at the site of future bouton formation or post bouton elimination were counted in the total contacts for each category of boutons.

**Figure 4.**
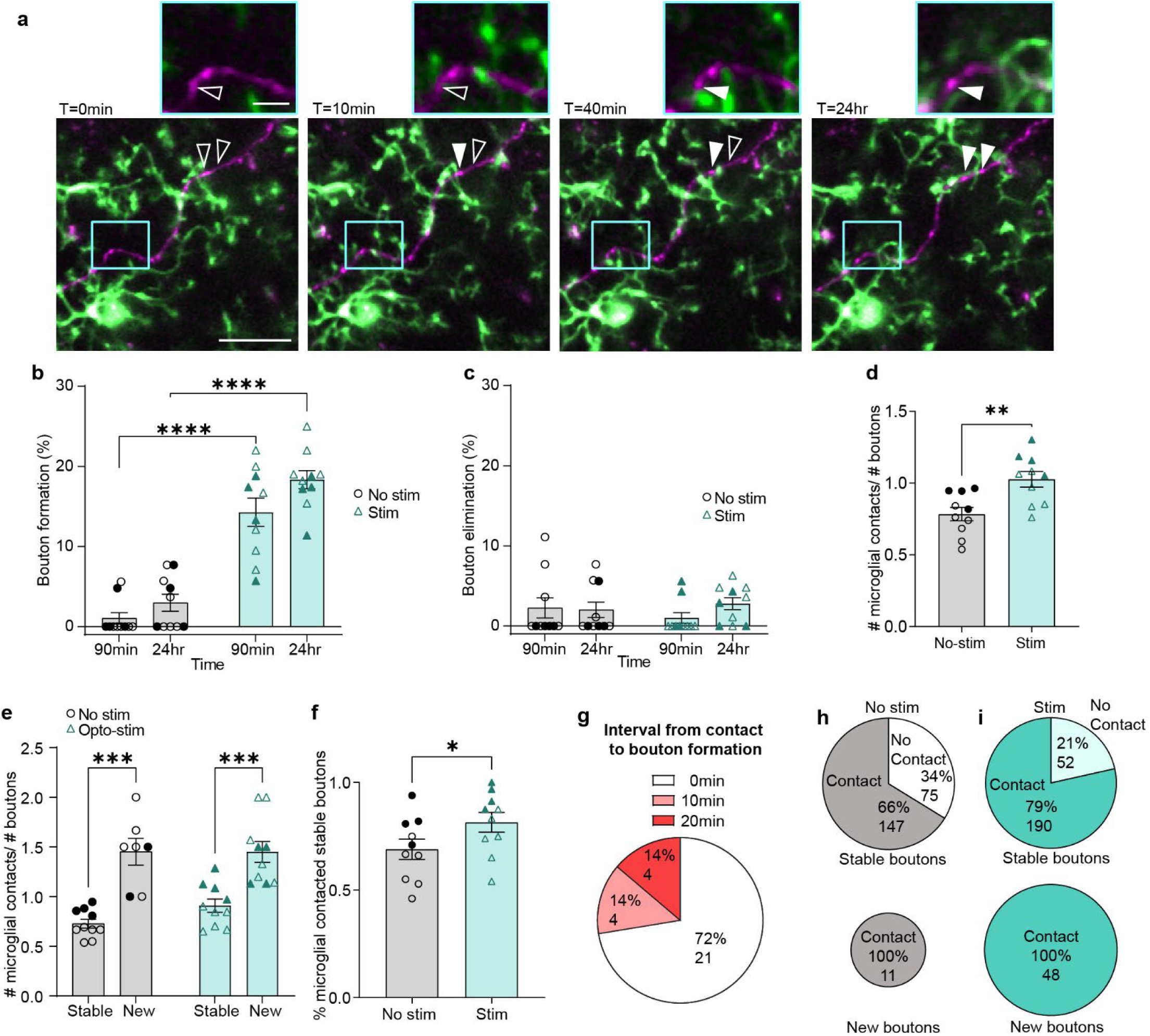
Microglia contact the axonal sites for newly formed boutons and increase surveillance of stable boutons after adolescent phasic DA axon activity. (**a**) Representative images of bouton formation with phasic DA axon stimulation (hollow arrowheads represent sites of future boutons, solid arrowheads denote a newly formed bouton). Rectangular pop-out highlights our observation that microglial processes contact axons prior to new bouton formation. (**b**) Phasic stimulation of DA axons drives bouton formation (n=10 mice, two-way repeated measures ANOVA stimulation p<0.0001, F(1,18)=128.6, Šídák’s multiple comparisons No stim v. Stim 90min p<0.0001, 24hr p<0.0001 (**c**) Phasic stimulation of DA axons does not drive bouton elimination (n=10 mice, two-way repeated measures ANOVA, stimulation p=0.8194, F(1,18)=0.05369, Šídák’s multiple comparisons test No stim v. Stim 90min p=0.5825, 24hr p=0.8121). (**d**) Microglia contact boutons more frequently after phasic DA axon stimulation (n=10 mice, two-tailed paired t-test, p=0.0049 t(9)=3.702).(**e**) Microglia make more frequent contacts with newly formed than stable boutons (n=10 mice, Mixed-effects model, Fixed effects [type III], Stability p<0.0001, F(1,9)=45.21, Šídák’s multiple comparisons Stable v. New No stim p=0.0001, Opto-stim p=0.0008). (**f**) After phasic DA axon stimulation, microglia contact a larger proportion of stable boutons (n=10 mice, two-tailed paired t-test, p=0.0329, t(9)=2.518). (**g**) Pie chart representation of new boutons formed within the 90min imaging period post-stimulation, which were sub-grouped by the time from microglial contact to bouton formation. 0min indicates that microglial processes were still in contact at the moment of formation. 2 boutons were excluded as they formed during the stimulation period and were not able to be assayed for contact. (**h,i**) Pie charts showing pooled percentage (%) and number of stable and new boutons contacted by microglia in no stimulation (**h**) and stimulated (**i**) conditions. A higher % of stable boutons are contacted by microglia after stimulation. New boutons are always contacted by microglia and are more abundant after stimulation (illustrated by the size difference of the pie charts). Scale bars 20µm. Graphs show mean ± S.E.M *p<0.05, **p<0.01, ***p<0.005, ****p<0.0001. No stim and stim experiments were conducted within the same animals. Individual points represent individual animals with females as hollow symbols and males as solid symbols.

Analysis of microglial contacts with DA boutons revealed a tight coupling between microglial dynamics and bouton plasticity. During the 90min post-stimulation imaging period, there was a significant increase in microglial contacts with DA boutons (Fig. 4d two-tailed paired t-test, p=0.0049, Supplemental video 5). When bouton contact data was subdivided based on bouton stability, we found that microglia make more contacts per new bouton than per stable bouton. (Fig. 4e Mixed-effects model, Fixed effects [type III], Stability p<0.0001). This pattern was present in both unstimulated and stimulated conditions; however, it is essential to note that there were far more new boutons in the stimulated (48 boutons) versus unstimulated condition (11 boutons). Thus, newly formed boutons contribute to the overall increase in microglial contacts per bouton after optogenetic stimulation. This suggests that optogenetic stimulation amplifies the natural recruitment of microglial processes to the sites of new bouton formation. In addition to the frequent contacts with the sites of new boutons, microglia contacted a larger proportion of the stable boutons post-stimulation (Fig. 4f, paired t-test, p<0.05). This suggests a specific change in the pattern of microglial interactions with stable boutons after optogenetic stimulation.

A final, and surprising finding is that microglial contact of the axon backbone precedes the formation of new boutons. In all cases where we could observe the axon backbone before new bouton formation, we found microglial processes contacting the axonal sites prior to the appearance of new boutons (Fig. 4a,g). In the case of the 29 new boutons which formed within the 90-min imaging period post-stimulation, 21 boutons first appeared while the microglial process was still in contact with the axon, whereas the rest appeared within 20 min after microglia contact (Fig. 4g). There were only 2 new boutons which formed within the 90min imaging period in the unstimulated condition, and these also received microglial contact prior to bouton formation. Another population of boutons were detected only at the 24-hr sampling time, so their temporal relationship with microglia contact could not be determined precisely due to the impracticality of imaging throughout the 24hr interval. Importantly, in both unstimulated and stimulated conditions, all new boutons were preceded by microglial axon backbone contact (Fig. 4h,i).

While there were few eliminated boutons in either condition (unstimulated 7 boutons and stimulated 8 boutons), we saw that like newly formed boutons, eliminated boutons received more frequent contacts than stable boutons and eliminated boutons were always contacted by microglial processes prior to elimination (Supplemental figure 5). We also confirmed that light stimulation in no-opsin control animals did not show any changes in bouton dynamics or shifts in microglial contacts with boutons (Supplemental figure 6). Taken together, our observations suggest that microglia are exquisitely sensitive to changes in DA activity, and there is a compelling connection between microglial contact and structural changes at the axon.

### Pharmacological manipulation of DA receptor signaling disrupts the biphasic microglial response to DA axon stimulation in adolescence

In order to further assess the role of DA signaling in the relationship between microglial contacts and DA bouton formation, we altered either D1 or D2 receptor function pharmacologically prior to our stimulation paradigm (Fig. 5a). Our previous work showed that administering the D2R agonist quinpirole (Quin 1mg/kg i.p.) was sufficient to block adolescent mesofrontal plasticity, implicating D2R stimulation as a negative regulator of mesofrontal plasticity^9^. Thus, we decided to test if this D2R manipulation would also produce changes in microglial dynamics. Additionally, because D1 and D2 receptors generate opposing G-protein coupled downstream signals^34^, we hypothesized that D1R signaling may then be a potential positive driver of mesofrontal plasticity and microglial dynamics. Thus, we used the D1R antagonist SCH23390 (SCH 1mg/kg i.p.) to reduce D1R activity and test its involvement in microglia dynamics.

**Figure 5.**
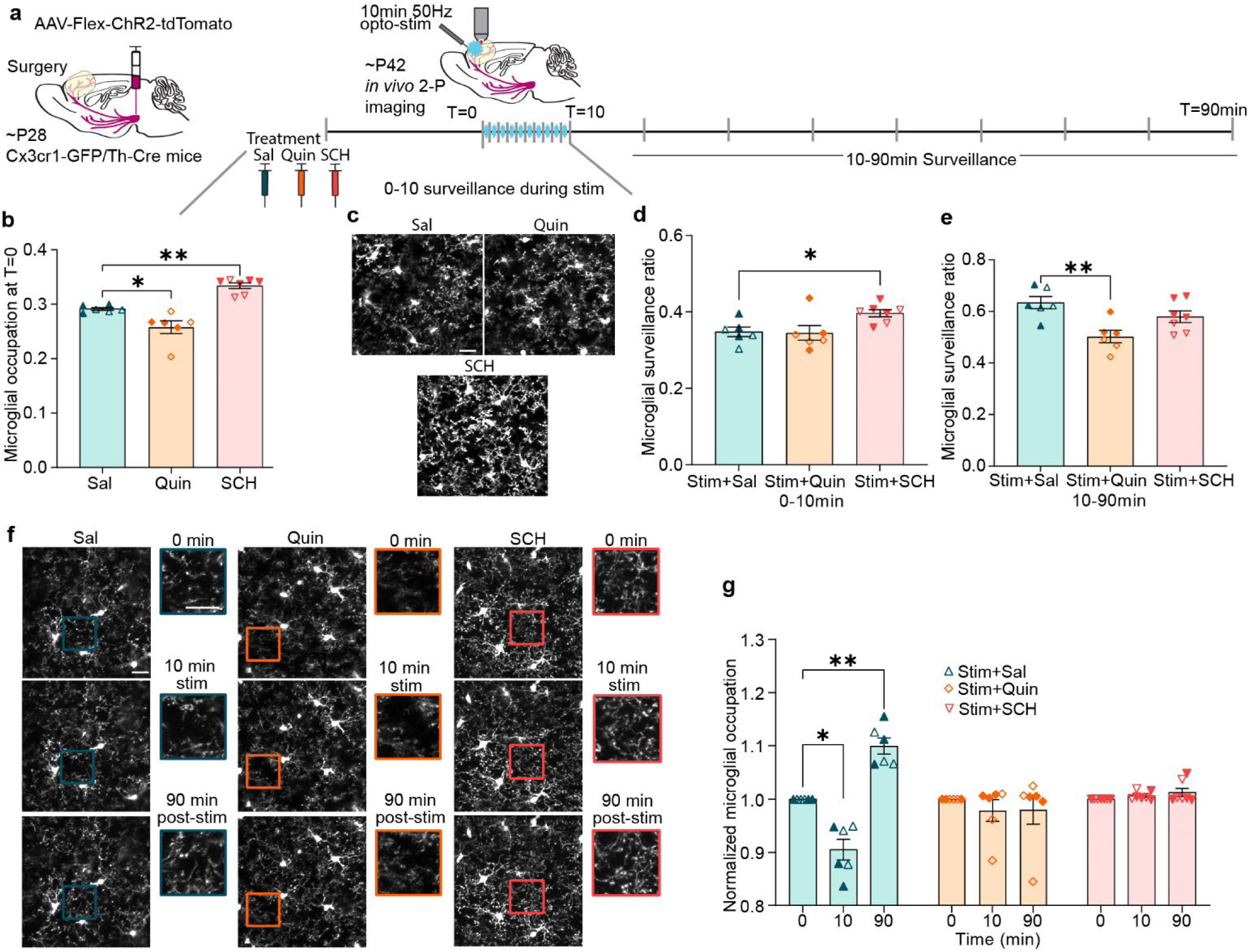
Manipulating DA receptor signaling disrupts adolescent microglial dynamics. (**a**) Timeline of dosing and imaging. (**b**) D2R agonism (Quinpirole 1mg/kg, i.p. Quin) decreases baseline microglial parenchyma occupation while D1R antagonism (SCH 23390 1mg/kg, i.p. SCH) increases baseline occupation (n: Sal=6, Quin=6, SCH=7 mice, one-way ANOVA p<0.0001 F(2,16)=28.21, Dunnett’s multiple comparisons test, Sal v. Quin p=0.0111, Sal v. SCH p=0.0013).(**c**) Representative images of microglial surveillance during phasic DA axon stimulation. (**d**) SCH prevents the expected decrease in microglial surveillance from 0-10min during DA axon stimulation (n: Sal=6, Quin=6, SCH=7 mice, one-way ANOVA p=0.0270, F(2,16)=4.565, Dunnett’s multiple comparisons Stim+Sal v. Stim+SCH p=0.0426). (**e**) Quin prevents increased surveillance from 10-90min post-DA axon stimulation (n: Sal=6, Quin=6, SCH=7 mice, one-way ANOVA p=0.0051, F(2,16)=7.490, Dunnett’s multiple comparisons test, Stim+Sal v. Stim+Quin p=0.0027). (**f**) Representative images of microglial occupation. Rectangle pop-outs denote magnified areas.(**g**) Both Quin and SCH prevent the dynamic changes in microglial occupation which accompany phasic DA axon stimulation (n: Sal=6, Quin=6, SCH=7 mice, two-way repeated measures ANOVA, Time x Treatment p<0.0001, F(4,32)=20.33, Tukey’s multiple comparisons Stim+Sal 0 v. 10min p=0.0103, 0 v. 90min p=0.0030, 10 v. 90 min p=0.0037). Scale bars 20µm. Graphs show mean ± S.E.M *p<0.05, **p<0.01, ***p<0.005, ****p<0.0001. Individual points represent individual animals with females as hollow symbols and males as solid symbols.

Interestingly, we found that administration of Quin significantly reduced baseline microglial occupation while SCH significantly increased occupation (Fig. 5b one-way ANOVA p<0.0001). From 0-10min during DA axon stimulation, microglia surveillance in the saline (Sal) control group decreased to a level similar to what we observed in our initial experiments without any saline or drug injection (Fig. 3c). The Quin treated group showed a similar level of surveillance to the Sal control, but the SCH treated group maintained a significantly higher level of surveillance and did not respond to DA axon stimulation (Fig. 5c,d one-way ANOVA p<0.05, Supplemental videos 6-8). In our initial optogenetic stimulation experiments, we also saw that 10-90min post-stimulation microglial surveillance increased (Fig. 3e). The Sal control group followed this pattern, achieving surveillance levels comparable to our first cohort (Fig. 5e). The SCH treated mice also had high microglial surveillance post-stimulation, but the Quin treated mice had decreased microglial surveillance (Fig. 5e one-way ANOVA p<0.01, Supplemental Videos 9-11). To better parse how Quin and SCH were impacting the biphasic microglial response to adolescent DA axon stimulation, we evaluated microglial occupation at 0, 10, and 90min. We found that like our previous stimulation cohort, the Sal treated control animals showed a reduction at 10min when stimulation had just ended and then a rebound in arborization by 90min post-stimulation. In comparison, there was no dynamic change in either Quin or SCH treated groups (Fig. 5f,g two-way repeated measures ANOVA, Time x Treatment p<0.0001).

Taken together, these results suggest that the pharmacological manipulation of D1R or D2R receptors was sufficient to cause a sustained change in microglial arborization and surveillance. D2R agonism induced microglial process retraction, rendering them in a state similar to that observed during DA axon stimulation, and precluded further dynamic surveillance changes in response to DA stimulation. On the other hand, D1R antagonism increased microglial arborization and kept them in this expanded state with enhanced surveillance, but precluded further dynamic responses to DA axon stimulation. Thus, D1R and D2R signaling may both be involved in the biphasic microglial response to DA axon stimulation.

### Altering DA receptor signaling disrupts adolescent mesofrontal axon plasticity and reduces microglial contacts with DA boutons

Given the literature highlighting the relationship between microglial arborization and their ability to interact with synaptic elements^14–18^, we next tested if the disruption in microglial dynamics from D1R and D2R receptor manipulation impacted how microglia interacted with DA boutons. We also investigated the potential concomitant changes in mesofrontal DA axon plasticity. We found that both D1R antagonism and D2R agonism blocked DA bouton formation after adolescent phasic DA axon stimulation (Fig. 6a Mixed-effects model, Fixed effects [type III], Treatment p<0.0001). Interestingly, at 24hr post-stimulation we also saw an increase in bouton elimination in our SCH group, suggesting that D1R antagonism may have had a destabilizing effect on DA bouton maintenance (Fig. 6b Mixed-effects model, Fixed effects [type III], Treatment p=0.0650, Dunnett’s multiple comparisons, 24hr Stim+Sal v. Stim+SCH p<0.05).

**Figure 6.**
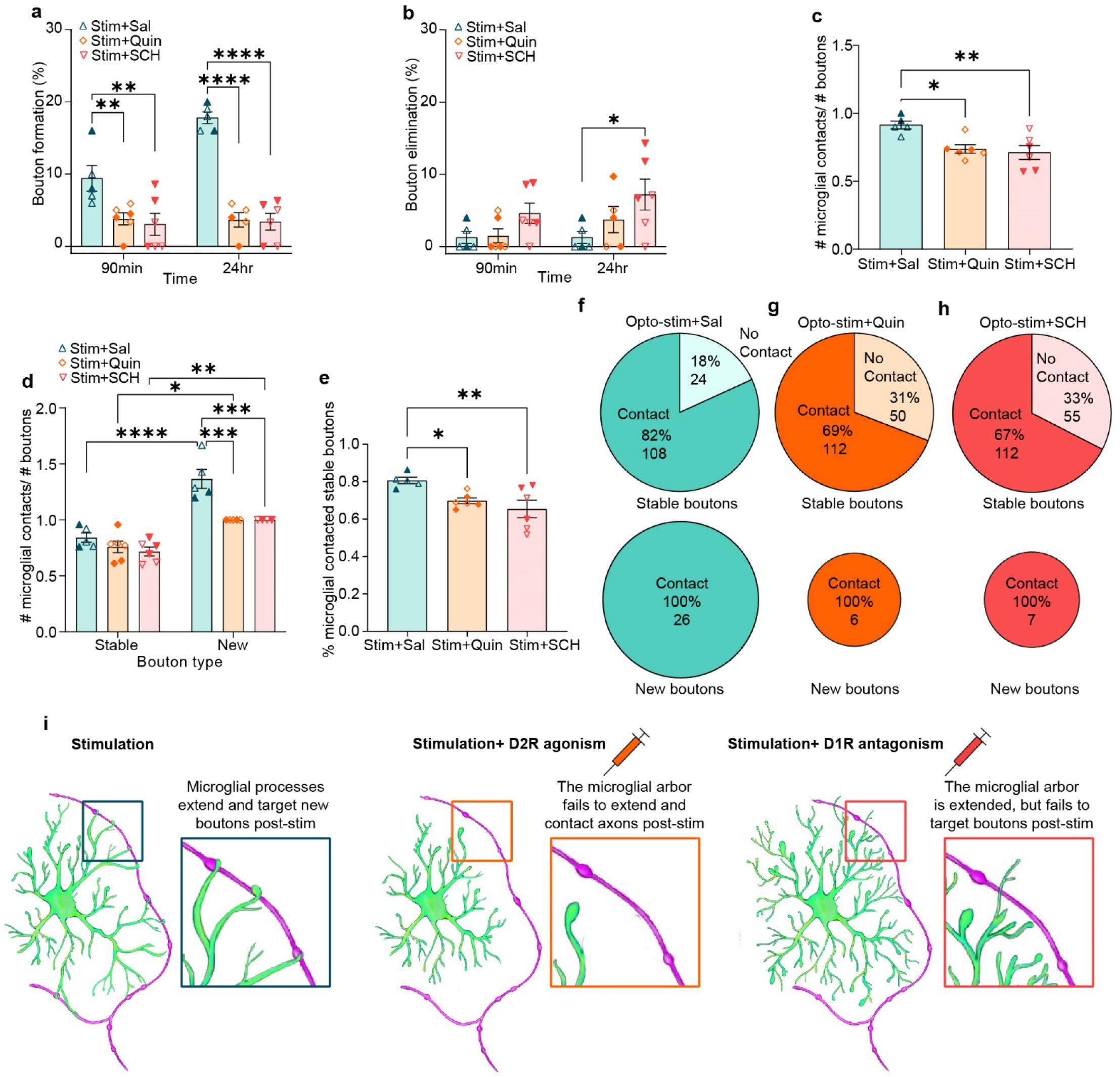
Manipulating DA receptor signaling blocks adolescent mesofrontal plasticity and reduces microglial contacts with DA boutons. (**a**) Both Quin and SCH blocked DA bouton formation post-phasic DA axon activation (n: Sal=5, Quin=6, SCH=6, Mixed-effects model, Fixed effects [type III], Treatment p<0.0001, F(2,14)=30.59, Dunnett’s multiple comparisons 90min: Stim+Sal v. Stim+Quin p=0.0065, Stim+Sal v. Stim+SCH p=0.0021, 24hr: Stim+Sal v. Stim+Quin p<0.0001, Stim+Sal v. Stim +SCH p<0.0001). (**b**) SCH increased DA bouton elimination 24 hr post-phasic DA axon activation (n: Sal=5, Quin=6, SCH=6 Mixed-effects model, Fixed effects [type III], Treatment p=0.0650, F(2,14) = 3.343, Dunnett’s multiple comparisons, 24hr Stim+Sal v. Stim+SCH p=0.0144). (**c**) Both Quin and SCH reduced the number of microglial contacts per bouton (n: Sal=5, Quin=6, SCH=6, One-way ANOVA p=0.0069, F(2,14)=7.264, Dunnett’s multiple comparisons test Stim+Sal v. Stim+Quin p=0.0144 Stim+Sal v. Stim+SCH p=0.0059). (**d**) While newly formed boutons still received more microglial contacts than stable boutons in all conditions, both Quin and SCH significantly reduced the number of contacts per new bouton compared to control mice (n: Sal=5, Quin=6, SCH=6, Two-way ANOVA Bouton category x Treatment p=0.0154, F(2,25)=4.951, Šídák’s multiple comparisons test Stable Stim+Sal v. New Stim+Sal p<0.0001 Stable Stim+Quin v. New Stim+Quin p=0.0189, Stable Stim+SCH v. New Stim+SCH p=0.0077, New Stim+Sal v. New Stim+Quin p=0.0003, New Stim+Sal v. New Stim+SCH p=0.0006). (**e**) Both Quin and SCH reduced the proportion of stable boutons that microglia contact (n: Sal=5, Quin=6, SCH=6, one-way ANOVA, p=0.0143, F(2,14)=5.839, Holm-Šídák’s multiple comparisons test, Stim+Sal v. Stim+Quin p=0.0306, Stim+Sal v. Stim+SCH p= 0.0096). (**f,g,h**) Pie charts showing pooled percentage (%) and number of stable and new boutons contacted by microglia in Sal (**f**), Quin (**g**), and SCH (**h**) conditions after opto-stimulation. Both Quin and SCH reduced the percentage of stable boutons contacted by microglia. New boutons were always contacted by microglia but were less abundant after Quin and SCH treatment (illustrated by the size difference of the pie charts). (**i**) Summary diagram of how DR perturbations alter microglial dynamics and interactions with boutons. Graphs show mean ± S.E.M *p<0.05, **p<0.01, ***p<0.005, ****p<0.0001. Individual points represent individual animals with females as hollow symbols and males as solid symbols.

We found that while the Sal control group showed a similar degree of microglia contacts per bouton as in the previous no-treatment experiment (Fig. 4d), both Quin and SCH groups showed significantly lower contacts compared to the Sal group, suggesting a failure to recruit microglial processes to boutons (Fig. 6c, One-way ANOVA, p=0.0069). While across all groups, microglia still contacted newly formed boutons more frequently than stable boutons, there was a significant reduction of this effect in both SCH and Quin groups (Fig. 6d Two-way ANOVA Bouton category x Treatment p<0.05). There also appeared to be more microglial contacts with eliminated boutons than stable boutons, but the number of eliminated boutons observed was low (Supplemental Figure 7). Furthermore, both Quin and SCH treated microglia interacted with a smaller percentage of stable boutons than that in the Sal control (Fig. 6e one-way ANOVA, p<0.05). Across the groups, microglia contacted all new boutons, but again there were far more new boutons in the Sal treated controls compared to Quin and SCH groups (Fig. 6f-h).

These microglia bouton interaction findings were especially surprising in the SCH group where the microglia occupied and surveyed the parenchyma at an increased rate comparable to that of the Sal stimulated control group. These findings suggest that increased microglial surveillance alone is not sufficient to produce increased microglia-DA bouton interactions. Our results indicate that microglial targeting to DA boutons is impaired when D1R receptor activity is blocked with SCH. Thus, our findings depict an interesting relationship between microglial dynamics and mesofrontal plasticity, whereby manipulating either D1R or D2R signaling alters microglial dynamics and blocks mesofrontal plasticity (Fig. 6i).

### Pharmacological inhibition of D2R in adults produces an “adolescent-like” microglial response to DA axon stimulation

After observing the robust microglial response to changes in DA signaling and mesofrontal plasticity during adolescence, we next tested if reinstating mesofrontal plasticity in adult animals would generate a similar microglial response. In adult mice, optogenetic stimulation of the VTA DA neurons alone does not generate mesofrontal DA plasticity, however pairing stimulation with a D2R antagonist reinstates plasticity^9^. Based on this study, we treated adult animals (∼P95) with eticlopride (Etic,1mg/kg, i.p.), a D2R antagonist, prior to our optogenetic stimulation paradigm (Fig. 7a). We found that Etic did not alter the baseline microglial occupation of the parenchyma (Fig. 7b unpaired t-test, p=0.5085), suggesting that any baseline activation of D2Rs from tonic release of DA likely had little impact on microglial dynamics.

**Figure 7.**
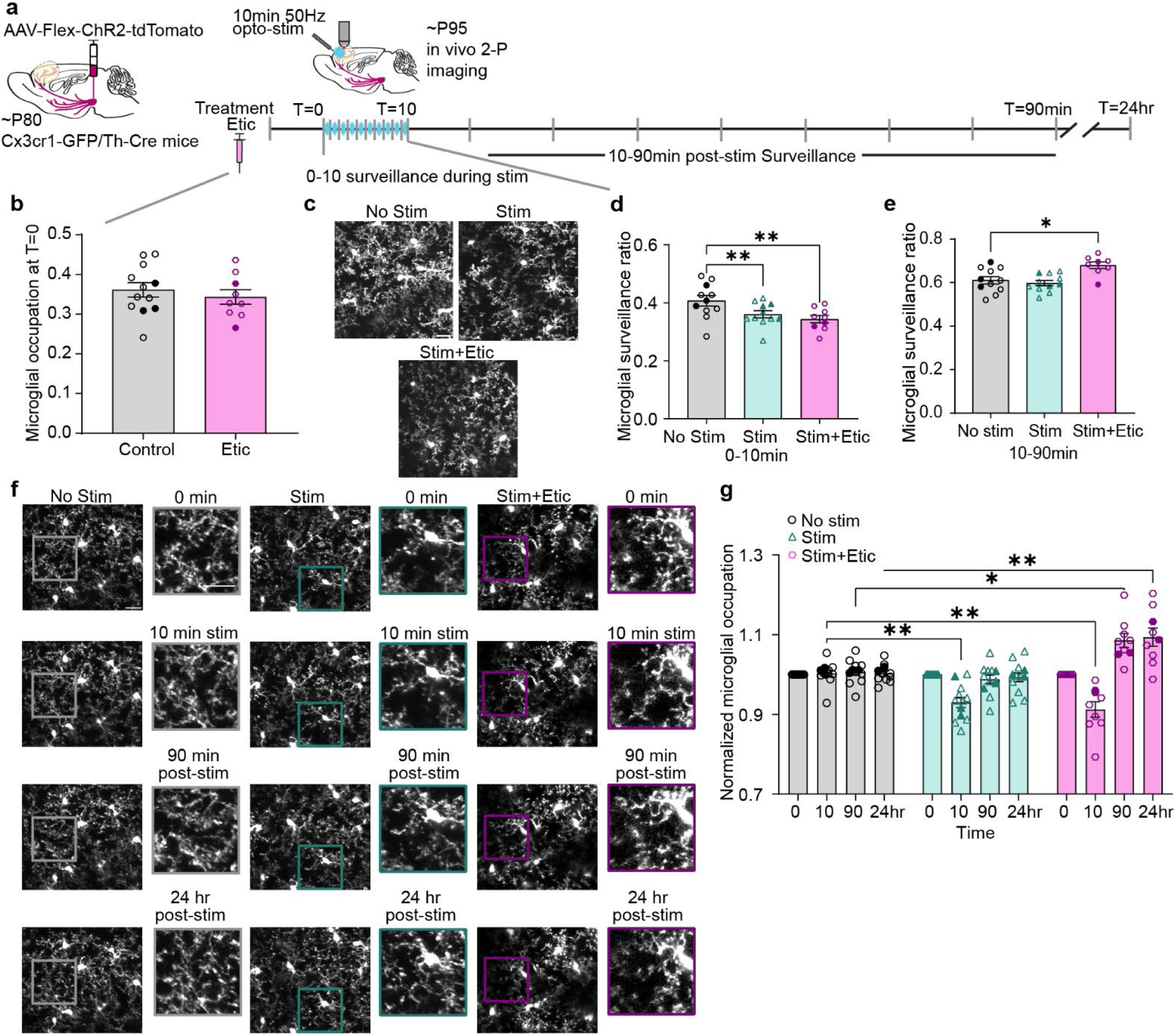
Inhibition of D2R produces an “adolescent-like” microglial response to DA axon stimulation. (**a**) Timeline of dosing and imaging. (**b**) D2R antagonism (eticlopride 1mg/kg, i.p. Etic) does not alter baseline parenchyma occupation (n: Control=12, Etic=9 mice, two-sided unpaired t-test t(19)=0.6738, p=0.5085).(**c**) Representative images of microglial surveillance during phasic DA axon stimulation. (**d**) Adult microglia respond to DA axon stimulation acutely independent of D2R inhibition (n: No stim=11, Stim=11, Stim+Etic=9 mice, Mixed-effects model, Fixed effect [Type III] p=0.003, F(1.827,16.44)=14.53, Dunnett’s multiple comparisons test, No Stim v. Stim p=0.0013, No Stim v. Stim+Etic p=0.0059). (**e**) D2R inhibition during stimulation restores post-stimulation increase in microglial surveillance in adults (n: No stim=11, Stim=11, Stim+Etic=8 mice, Mixed-effects model, Fixed effect [Type III] p=0.0029, F(3.126,16.28)= 8.696, Dunnett’s multiple comparisons test, No Stim v. Stim p=0.7707 No Stim v. Stim+Etic p=0.0211). (**f**) Representative images of microglial occupation (Pop-outs highlight areas of interest). (**g**) D2R inhibition during stimulation restores the biphasic microglial response to mesofrontal activity (n: No stim=11, Stim=11, Stim+Etic=8 mice, Mixed-effects model, Fixed effect [Type III] Treatment x Time p<0.0001, F(3.126,27.61)=16.36, Dunnett’s multiple comparisons test, 10min: No Stim v. Stim p=0.0020, No Stim v. Stim+Etic p=0.0022, 90min: No Stim v. Stim+Etic p=0.0163, 24hr: No Stim v. Stim+Etic p=0.0017). Scale bars 20µm. Graphs show mean ± S.E.M *p<0.05, **p<0.01, ***p<0.005, ****p<0.0001. No stim, stim and stim+etic experiments were conducted within the same animals. Individual points represent individual animals with females as hollow symbols and males as solid symbols.

Next, we evaluated the response of adult microglia to mesofrontal DA axon stimulation. Like adolescent microglia, adult microglia acutely responded to phasic DA stimulation with a reduction in surveillance. This effect was independent of D2R inhibition (Fig. 7c,d, Mixed-effects model, Fixed effect [Type III] p=0.0029, Supplemental Videos 12-14), suggesting that microglial retraction in response to DA stimulation may be mediated through other receptors or secondary signals. The impact of D2R inhibition became apparent in the post-stimulation period, as Etic restored the post-stimulation increase in microglial surveillance in adults (Fig. 7e, Fixed effect [Type III] p=0.003, Supplemental Videos 15-17). Unlike the DA axon stimulation alone condition, stimulation paired with Etic generated the full biphasic microglial response that we had observed in our adolescent animals (Fig. 7f,g, Mixed-effects model, Fixed effect [Type III] Treatment x Time, p<0.0001). Taken together, these results suggest that pharmacological inhibition of D2R in adults produces an “adolescent-like” microglial response to phasic DA axon stimulation.

### D2R inhibition in adults restores both mesofrontal axonal plasticity and microglial recruitment to DA boutons

After observing a biphasic response in adult microglia with Etic administration, we next examined the effects of D2R inhibition on microglial interactions with DA boutons. First, we confirmed that as in our previous work, D2R inhibition during DA axon stimulation reinstates plasticity in adults, driving bouton formation without a concomitant change in elimination (Fig. 8a,b Formation Mixed-effects model, Fixed effects [type III], Treatment p<0.0001; Elimination Mixed-effects model, Fixed effects [type III], Treatment p=0.6992). Importantly, we found that whereas DA axon stimulation alone did not promote microglial contacts with boutons, the addition of Etic increased the number of microglial contacts per bouton (Fig. 8c Mixed-effects model, Fixed effects [type III], p=0.0002). This result implies that microglia are actively recruited when the circuit is being remodeled, rather than just in response to changes in neural activity.

**Figure 8.**
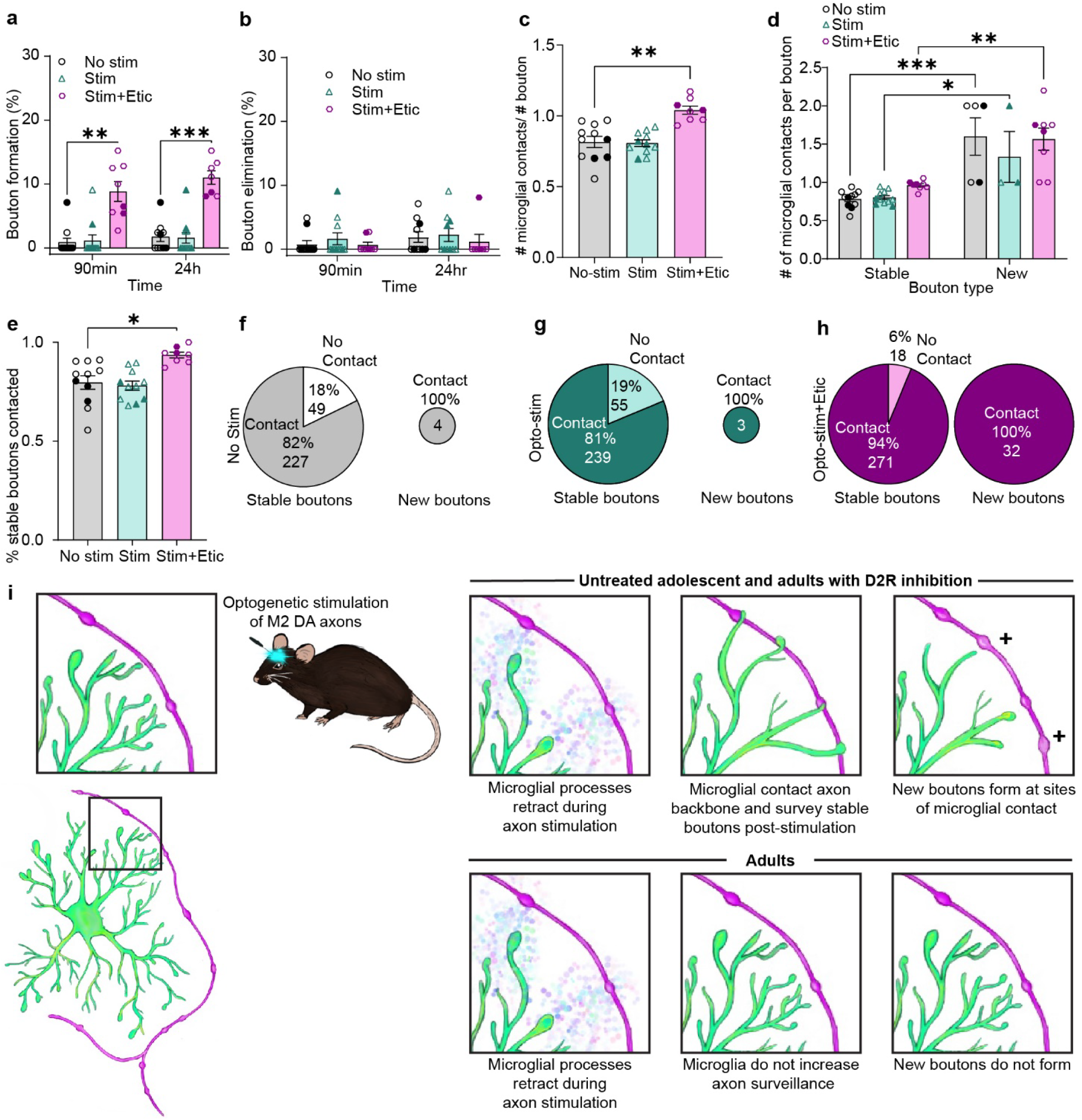
D2R inhibition restores both mesofrontal axonal plasticity and microglial recruitment to DA boutons in adults. (**a**) D2R inhibition during DA axon stimulation reopens adult plasticity (n: No stim=11, Stim=11, Stim+Etic=8, Mixed-effects model, Fixed effects [type III], Treatment p<0.0001, F(1.110,11.10)=51.27, Dunnett’s multiple comparisons 90min: No stim v. Stim+Etic p=0.0017, 24hr: No Stim v. Stim+Etic p=0.003). (**b**) Etic does not alter bouton elimination (n: No stim=11, Stim=11, Stim+Etic=8, Mixed-effects model, Fixed effects [type III], Treatment p=0.6992, F(2,27)=.3626). (**c**) Etic increases the number of microglial contacts with boutons after DA axon stimulation (Mixed-effects model, Fixed effects [type III], p=0.0002 F(1.516,12.89)=19.92, Dunnett’s multiple comparisons No Stim v. Stim+Etic p=0.0019) (**d**) Microglia preferentially make more contacts with new boutons than stable boutons (Mixed-effects model, Fixed effects [type III] bouton type p<0.0001 F(1,13)=51.59, Šídák’s multiple comparisons Stable v. New: No stim p=0.0003, Stim p=0.0338, Stim+Etic p=0.0021). (**e**) Etic increases the proportion of stable boutons surveyed by microglia after DA axon stimulation (Mixed-effects model, Fixed effects [type III] p=0.0042 F(1.518,12.90)=9.812, Dunnett’s multiple comparisons No stim v. Stim+Etic p=0.0107). (**f,g,h**) Pie charts showing pooled percentage (%) and number of stable and new boutons contacted by microglia in no-stimulation (**f**), stimulation (**g**), and stimulation + Etic (**h**) conditions. Etic increased the percentage of stable boutons contacted by microglia. New boutons were always contacted by microglia but were more abundant after Etic treatment (illustrated by the size difference of the pie charts). (**i**) Summary diagram of how D2R inhibition in adults restores the “adolescent-like” relationship between microglia and DA boutons. Graphs show mean ± S.E.M *p<0.05, **p<0.01, ***p<0.005, ****p<0.0001. No stim, Stim and Stim+Etic experiments were conducted within the same animals. Individual points represent individual animals with females as hollow symbols and males as solid symbols.

Like in adolescent animals, microglia in adults preferentially made more contacts with new boutons compared to stable boutons, although few new boutons appeared in adults unless stimulation was paired with Etic (Fig. 8d Mixed-effects model, Fixed effects [type III] p<0.0001). There were also more microglial contacts with eliminated boutons than stable boutons, but the number of eliminated boutons observed was low (Supplemental Figure 8). Importantly, Etic increased the proportion of stable boutons surveyed by microglia, again paralleling our stimulated adolescent animals (Fig. 8e Mixed-effects model, Fixed effects [type III] p=0.0042). As in our adolescent cohorts, we found all new boutons were contacted by microglia, however, again there were significantly more new boutons in the stimulated animals that received D2R inhibition than the unstimulated and stimulation alone groups (Fig 8f-h). Thus, inhibiting D2R signaling in adult mice reinstates mesofrontal axonal plasticity with the recruitment of microglial processes to DA boutons (Fig. 8i). Taken together, our data from both adolescent and adult animals highlight the intimate relationship between microglial surveillance of axons and DA bouton formation during mesofrontal plasticity.

## Discussion

We report, for the first time, that frontal cortical microglia are exquisitely sensitive to DA signaling, and microglial surveillance is engaged in adolescent mesofrontal plasticity. Our study integrated a range of advanced technical approaches, including dual genetic labeling of microglia and DA projections, longitudinal *in vivo* two-photon imaging of glial-neural interactions in awake animals, ethologically relevant behavior paradigms, projection-specific optogenetic manipulation, and complementary pharmacological interventions. We found that adolescent frontal cortical microglia exhibit a unique biphasic response to DA axon stimulation with an initial retraction and a subsequent extension of their processes. This robust process extension facilitated increased microglial surveillance of DA boutons. Critically, microglial contacts along the axon backbone preceded new bouton formation. Perturbation of either D1 or D2 signaling in adolescence was sufficient to impair both the biphasic microglial response and mesofrontal plasticity. Finally, we found that inhibiting D2 signaling in adulthood produced an “adolescent-like” microglial response to phasic DA stimulation and reinstated mesofrontal plasticity. Our findings highlight an intimate relationship between microglial surveillance and activity dependent DA bouton formation. Furthermore, our results suggest that D1 and D2 signaling are central regulators of microglia-DA axon interactions and adolescent mesofrontal plasticity.

### Microglial surveillance and DA bouton formation in the frontal cortex

The advent of *in vivo* two photon microscopy techniques enabled new insights into the non-pathological dynamics of microglial surveillance^35,36^ and the interactions of dynamic microglial processes with neuronal elements in the intact CNS^37,38^. Since the initial discovery of microglial surveillance of neural elements, extensive work has shown that microglia are essential participants in developmental plasticity^14,39,40^. Much of the previous *in vivo* and *ex vivo* work has identified microglia as facilitators of dendritic spine or synaptic bouton removal during periods of development or network remodeling^38,41,42^. Microglial immune machinery such as the complement system is frequently repurposed for synaptic pruning during health and disease^14,43^. As immune cells, microglia are naturally suited to removing CNS elements, however our work highlights microglia acting to facilitate growth rather than elimination of neuronal structures.

In the context of the existing literature, our present study provides several novel insights into microglial-neuron dynamics. First, to our knowledge, this is the first *in vivo* description of microglial interactions with neuromodulatory boutons, which are not associated with well-defined postsynaptic structures like glutamatergic or GABAergic boutons^11^. Second, this is the first study associating microglial contact with axonal bouton formation rather than removal^38^. We identified many instances of tight temporal coupling between microglial contact and bouton formation post axon stimulation, indicating that the rapid recruitment of microglial processes to the axon backbone may be an essential step in bouton formation. Our observation that DA bouton formation is always preceded by microglial contact of the axon backbone suggests that microglial processes may be remodeling the extracellular matrix^44^ or triggering axonal reorganization to facilitate new bouton formation.

Based on the previous work, there are multiple candidate molecular mechanisms by which microglia could facilitate bouton formation. Microglia have a wide repertoire of receptors and signaling systems which allow them to guide plasticity and development^41^. In the hippocampus, neuronal interleukin 33 signaling drives microglial extracellular matrix remodeling necessary for hippocampal dendritic spine and synapse formation^44^. In the developing somatosensory cortex, microglial contacts with dendrites were associated with calcium transients and actin accumulation preceding the formation of filopodia^45^. In the motor cortex, microglia were found to facilitate dendritic spine formation during motor learning through release of brain derived neurotropic factor^17^. However, these examples suggest that the mechanisms by which microglia guide synaptic remodeling and plasticity are not uniform across regions of the CNS^41^. The precise molecular mechanisms connecting microglial surveillance to DA bouton formation in the frontal cortex present exciting directions for future research. Although the lack of a uniform mechanism of microglia-neuron interactions complicates our understanding of how microglia impact neuronal plasticity, it also means that manipulations of microglial roles in development and plasticity can be highly targeted and specific. We found that wheel running generated a regionally specific microglial response, highlighting the possibility to selectively target frontal cortical microglia with a naturalistic rewarding behavior.

### Unique responses of frontal cortical microglia to DA signaling in adolescence

Adolescence is increasingly being viewed as a sensitive period for the development of the frontal cortex^1^. Microglia have been implicated as key regulators of sensitive developmental periods across multiple cortical areas, including the frontal cortex^16,46^. As immune cells that are acutely sensitive to changes in the CNS milieu, microglia are uniquely suited for mediating rapid and specific changes in neural circuitry. However, there are many outstanding questions, not only on how microglia interface with neuronal elements, but also what signals regulate microglial participation during sensitive periods of development.

Our study provides several lines of evidence indicating that frontal cortical microglia respond uniquely to DA signaling during adolescence. First, using voluntary wheel running as a naturally rewarding behavioral paradigm for mice, we found that this rewarding experience promotes microglial process outgrowth specifically in the adolescent frontal cortex, which receives extensive DA innervation, but not in the adolescent visual cortex, which receives little DA innervation. Second, by directly stimulating frontal DA axons with a phasic activity pattern typically associated with reward, we discovered a biphasic microglia response characterized by a transient arbor retraction during DA stimulation and a sustained arbor extension post-stimulation. Third, we found that D1 antagonism keeps microglia arbors in an extended state, whereas D2 agonism keeps microglia arbors in a retracted state, both preventing biphasic microglial responses to frontal DA input. While previous studies have identified that microglia surveillance of the parenchyma can be inhibited by noradrenergic signaling^15,21^, this biphasic microglial response has not been reported before. Furthermore, the sustained increase of microglial surveillance after rewarding experience or DA axon stimulation is another new finding of this study.

DA could be directly influencing microglial physiology, as previous work has found microglia express both D1 and D2-type DA receptors (DRs)^24,47^ and that signaling through DRs impacts microglial membrane conductance, chemotaxis, phagocytosis, and interleukin signaling *in vitro*^24,48,49^. However, previous studies did not provide *in vivo* evidence of whether or how microglia respond to DA signaling. Interestingly, only a subset of microglia in cell culture responded to DA with a change in inward rectifying K_ir_ conductance, underscoring the potential cellular heterogeneity^24^. Previous work has also highlighted the importance of the K^+^ channel THIK-1 in maintaining the resting membrane potential of microglia and resting microglial surveillance of the CNS^50^. The *in vitro* effects of DR signaling on microglial membrane conductance^24^ raise the possibility that DRs may interface with THIK-1 or K_ir_ K^+^ channels to alter microglial surveillance of the parenchyma.

Furthermore, it is also conceivable that DA may influence microglial dynamics through other cell types and molecular pathways. Both excitatory and inhibitory frontal cortical neurons express DA receptors and respond to DA signaling^5,34^. Microglia can sense changes in fast neurotransmission as well as activity-dependent local release of ATP^41^. Recent work has found that microglial protrusions are recruited to thalamocortical synaptic boutons in an activity dependent manner by purinergic signaling through microglial P2RY12 receptor^51^. We show that microglia have a biphasic response to DA signaling, with an initial process retraction followed by extension. D1 inhibition leads to extended microglial processes at the basal state and prevents the biphasic response to subsequent DA stimulation. In addition, microglial recruitment to DA boutons is disrupted, despite the extended microglial processes. Thus, D1 signaling may drive the initial microglial retraction and prime microglial processes for subsequent re-targeting, while activity-dependent co-release of ATP and purinergic signaling may facilitate process chemotaxis towards activated DA axons^51,52^. Additionally, astrocytes also express DRs^53^, and thus could be another mediator of the microglial response to mesofrontal DA signals^54^. It will be an interesting research direction to dissect how neuronal, astroglial, and/or microglial DR signaling may mediate the microglial response to DA.

### Therapeutic implications of microglial interactions with DA circuits

Abnormalities in adolescent DA system development have long been associated with neurodevelopmental disorders, such as schizophrenia^55,56^. Recently, microglial dysfunction has increasingly been linked to neurodevelopmental disorder pathology^57^. In pathological conditions, microglia immune reactivity can result in their failure to execute their homeostatic roles^41^ and they can also drive unwanted synapse pruning^43,58^. Here, we report a compelling relationship between microglial contact with axons and subsequent activity-dependent DA bouton formation. D2 agonism in adolescence suppresses microglial surveillance and bouton formation, whereas D2 antagonism in adults reinstates an adolescent-like microglial surveillance and mesofrontal plasticity. Interestingly, D2 inhibition does not affect the basal state of microglia processes in the adult frontal cortex or the initial retraction during DA stimulation, but restores the increased extension and bouton contact after stimulation, which is otherwise only observed in adolescence. This result suggests that age-dependent changes in D2 signaling^5^ may constrain microglia surveillance and reduce activity-dependent plasticity in the adult frontal cortex.

While the exact cell type and spatial location where a systematically administered D2 antagonist acts to promote frontal cortical microglial surveillance and neural plasticity remains open for further investigation, our study provides an interesting animal model to investigate the therapeutic potential of systematically delivered antipsychotic medicines. Because antipsychotic medicines are known to engage D2 antagonism^7^, our findings raise the possibility that a combination of pharmacological antipsychotic treatment and behavioral DA stimulation, such as through exercise^59^, could be used to remedy frontal DA deficiencies in psychiatric disorders by promoting microglia surveillance and activity-dependent DA bouton outgrowth. In summary, our study provides important evidence that DA signaling regulates microglial surveillance and adolescent plasticity in the frontal cortex and lays the groundwork for future investigations into the therapeutic implications of microglial interactions with DA circuits in health and disease.

## Acknowledgements

The authors thank A. Majewska, Z. He and members of the Wang lab for critical discussion and reading of the manuscript, and L. Shaw for technical advice. This work was supported by grants from National Institutes of Health (R01MH127737 to K.H.W., F32MH124298 to R.S.) and Del Monte Institute for Neuroscience at the University of Rochester (to K.H.W.)

## Author Contributions

R.D.S. and K.H.W. conceived the study and designed the experiments. R.D.S. conducted the experiments and analyzed the data. R.D.S. and K.H.W. wrote the manuscript.

## Methods

### Animals

All experimental protocols were approved by the Institutional Animal Care and Use Committee at the University of Rochester and followed the National Institutes of Health guidelines. All experiments were conducted on both male and female mice on a C57BL/6 background between P28-P52 (adolescent) and P70-P100 (adult). CX3CR1^GFP^ (JAX 005582)^1^ were crossed to the TH-Cre^2^ line to allow for both microglial visualization and specific viral targeting for labeling and manipulation of dopaminergic axons. Mice were housed under a standard 12hr light/ 12hr dark cycle and fed standard chow and water ad libitum. In all figures, male mice are solid symbols and females are hollow, the numbers of each are also included in the figure legends.

### Pharmacologic agents

Quinpirole, a D2 receptor agonist, (Sigma, 1mg/kg), SCH23390, a D1 receptor antagonist, (EMD Millipore,1mg/kg), and eticlopride, a D2 receptor antagonist (1mg/kg) were used to acutely alter dopaminergic receptor signaling during *in vivo* optogenetic imaging experiments. Solutions were prepared fresh on the day of imaging using sterile saline. Each of these agents were injected i.p. ∼15min prior to phasic optogenetic stimulation in their respective experiments.

### Cranial window surgery and viral targeting of VTA dopaminergic neurons

Mice were kept under isoflurane anesthesia (2-3%) for the duration of the surgical procedures and a heating pad was used to maintain body temperature at 37°C during surgery. Mice were mounted and head fixed in a stereotaxic frame for all surgical procedures. All surgical procedures adhered to aseptic technique. After a scalp incision and clearing of the connective tissue, a 0.5-mm drill bit (FST) was used to drill a small hole through the skull for injection into the VTA. Either 375nl AAV9-CAG-FLEX-tdTomato (wheel running and optogenetic control, Addgene 28306-AAV9, 3.8 x 10^13^ gc/mL) or AAV9-CAG-Flex-ChR2-tdTomato (optogenetic stimulation, Boston Children’s Hospital Viral Core, 4.82 x 10^13^ gc/mL) was injected through a glass micropipette connected to a micropump (from bregma: AP -3.2, ML 0.5, and DV 4.5 mm, for both adolescent and adult experiments). After the VTA injection in the same surgical session, a 3mm craniotomy was opened over M2 using a dental drill. As previously described^3^, a 5mm coverslip attached to a 3mm coverslip (Electron microscopy sciences) by UV glue (Norland Optical Adhesive, Norland) was fixed into the craniotomy using dental cement (C&B Metabond, Parkell). Dental cement was then used to cover the remaining surface of the skull, seal the incision site, and affix a small metal head plate. Animals were given slow-release buprenex (5mg/kg subcutaneously every 72hrs) and monitored daily during the 72hr postoperative period. Animals were imaged no sooner than 2 weeks post-op to allow for full surgical recovery and viral construct expression. After the completion of all imaging experiments, all animals were perfused to verify viral labeling of the VTA. Any animals lacking viral labeling of the VTA were excluded from further analysis.

### Two-photon microscopy

A two-photon microscope (FV1000, Olympus) was used for all *in vivo* imaging (excitation laser: 900nm). All imaging was collected with a x25 water immersion lens (NA1.05) in head fixed awake animals with a 380-560 filter for GFP labeled microglia and a 575-630 filter for tdTomato labeled dopaminergic axons. Mice received three consecutive days of head restraint training prior to *in vivo* imaging sessions, progressing from 30 min to 90 min over the course of the training sessions. A 1x image was taken at the first time point for each animal in the experiment to facilitate locating the same ROI again at subsequent time points. For microglial occupancy and dopaminergic bouton analysis in wheel running experiments, individual z stacks of the same ROI were taken at 3x digital zoom with a 1-µm z step at each time point. For optogenetic experiments, repeated z stack imaging was done at an interval of every minute for 10 minutes (during stimulation) and every 10 minutes for 90 minutes (post-stimulation). Further details on the separate imaging parameters for these experiments are detailed below. All image analysis was done offline using ImageJ blind to condition as described in Stowell et al.^3^ and as described below.

#### Wheel Running

An initial baseline z stack ROI (100-150 µm, 1-µm z step) containing both labeled axons and microglia was collected, and then the animal was returned to its home cage, or placed on a running wheel (Med Associates) for 2hrs^4^. Med Associates Software was used to automatically record the number of revolutions the animal ran on the wheel. Any animals that ran fewer than 250 revolutions were excluded from further analysis. Immediately after the 2hr period, the ROI was imaged again, and then it was imaged a final time at 24hr. Images from the wheel running experiment were analyzed offline using ImageJ for bouton dynamics and microglial occupation, as described in the respective sections below.

#### Optogenetic stimulation

For stimulation experiments, two initial ROIs were located and imaged at baseline prior to stimulation. The ROI1 z-stack was used to observe the acute microglial response to DA axon stimulation between stimulation pulses (20 µm, 1-µm z step) and the ROI2 stack was used for the microglial bouton interaction measurements (60-80 µm, 1-µm z step. For optogenetic stimulation, an optical fiber (200 µm diameter, Thor Laboratories) was connected to a 473nm solid-state laser diode (CrystaLaser) with ∼20mW output from the fiber. This optical fiber was then positioned directly over the M2 cranial window glass coverslip to deliver phasic stimulation (50Hz pulse train, 3ms/pulse, 10 pulses/train, 1 train/min for 10 min). To measure the acute microglial response to DA axon stimulation, microglia in ROI1 were imaged every minute between the pulses for the 10-min stimulation period (10 stimulation z-stacks). To image microglial interactions with boutons and bouton dynamics, after the cessation of stimulation, ROI2 was imaged at 10min intervals for 90mins post stimulation (9 post-stimulation z-stacks). ROI2 was also imaged again at 24hrs for a final time point. Images from the optogenetic experiments were analyzed offline using ImageJ for bouton dynamics, microglial contacts with boutons, microglial occupation, and microglial surveillance as described in the respective sections below.

#### Calcium imaging

To verify that the optogenetic stimulation paradigm was effectively stimulating the mesofrontal circuit, we used *in vivo* two-photon calcium imaging to capture the cortical response to our stimulation paradigm. We used Th-cre mice and injected 375nl AAV2/9-CAG-Flex-ChR2-tdTomato into the VTA and infused 450nl AAV9.CamKII.GCaMP6s.WPRE.SV40 (Addgene 107790-AAV9, 2.5 x 10^13 GC/mL) into M2 (Bregma, AP 1.7, ML 0.8) from the pial surface using a glass micropipette connected to a micro-syringe pump as previously described^5^. A cranial window and head plate were affixed to the skull as described in the above surgical procedures. Optogenetic stimulation was done following the same phasic parameters. During stimulation, a time series of images (∼40s, 0.351s/frame) were collected between each stimulus train for a total of 10 time series between the pulses. Analysis was conducted offline using custom MatLab scripts. Briefly, baseline fluorescence (F0) was defined as the average fluorescent signal (Ft) in the first 15s of the time series. Changes in the calcium signaling (ΔF/F) was calculated as (Ft-F0)/F0. We compared the average of the 10 time series to the baseline calcium activity in the ROI.

#### Microglial Occupation and Surveillance

All analysis was done blind to condition. For all microglial analysis, the z stacks collected during each imaging session were processed in ImageJ to register any drifts over time, and remove movement artifact from animals moving during imaging (Correct 3-D drift, ImageJ). Time course images were also corrected for intensity shifts associated with photo bleaching (Bleach correct, ImageJ). The same section of the z stack was maximum intensity projected at each time point in an imaging series (20-30µm thick) and lateral motion artifacts were corrected (Stackreg plugin, ImageJ). To quantify microglial occupation at each timepoint in an image series, the z projections were binarized (Threshold, ImageJ) and the ratio of microglia occupied pixels out of the total ROI pixels was found for each time point. All occupation values were then normalized to the microglial occupation at the t=0 image. Surveillance analysis was performed as previously described^3^. Briefly, for surveillance analysis, consecutive z projections from each time point were collapsed through the maximum-intensity z-projection function in ImageJ. This image was binarized and surveillance was measured as the ratio of microglia-occupied pixels out of the total pixels in the image.

#### Bouton formation and elimination

To quantify bouton dynamics in all time-lapse *in vivo* two-photon images, boutons at each time point were labeled with the multi-point tool in ImageJ. Boutons were identified as bright swellings along axons larger than 0.5 µm^2^ in size with a fluorescent intensity 2.5-fold brighter than the adjacent axon backbone, as in previous work^4^. Bouton identification in each image was done blind to the condition. The percent bouton formation was calculated as the number of boutons present at time point 2 (either 90min, 2hr, or 24hr), but not time point 1 (0min) divided by the total number of boutons present at time point 1. The percentage of eliminated boutons was calculated as the number of boutons present at time point 1 but not at time point 2, divided by the total number of boutons present at time point 1.

#### Microglial bouton contact analysis

All contact analysis was performed blind to condition. To visualize microglial contacts with dopaminergic axons, the GFP (microglia channel) was assigned the color green and the tdTomato (axon channel) was assigned the color magenta. Microglial contacts with boutons or the axon backbone were determined by manually stepping through the z stack (60-80 µm, 1-µm z step) of the merged channels. A contact was counted as the colocalization of microglial and axonal pixels. For each animal, 10-50 boutons were evaluated for microglial contact per imaging condition collected (only axons visible at all time points were evaluated). While viral labeling efficacy varied between animals, within the same animal, similar numbers of boutons were counted in each condition. Animals with a higher number of labeled boutons had a greater number of boutons analyzed for contact in both unstimulated and stimulated conditions and vice versa for less robustly labeled animals. The multi-point tool in ImageJ was used to label and number boutons for time course tracking. All identifiable boutons were tracked through each time point to establish if it was stable, newly formed or eliminated. The total number of microglial contacts, whether the contact preceded a bouton change, and the interval from contact to bouton formation, were recorded. The number of contacts per bouton was calculated to determine the frequency of microglial bouton contacts. The proportion of boutons contacted was used to assess what percent of the total boutons in a given ROI were contacted by microglia. The contact frequency and proportion contacted data were subdivided by the categories of stable bouton, newly formed bouton, or eliminated bouton. For newly formed boutons, contacts with the axon back bone at the site of the new bouton prior to bouton formation were logged as bouton contacts and counted in the total count. To avoid issues of photo bleaching from repeated imaging, unique ROIs were selected for repeated measures within the same experimental animal.

### Histology

Intact brains were collected after transcardial perfusion with PBS and fixed overnight in paraformaldehyde (4%). Coronal sections of tissue were cut on a vibratome at 70-µm thickness. Sections were then mounted and cover slipped. For confirmation of viral injection efficacy, the VTA was viewed at 10x and 25x magnification using a confocal microscope (Olympus FV 1000). We also assessed the utility of the CX3CR1^GFP^/DAT-Cre (JAX: 006660)^6^/Ai14 (JAX:007914)^7^ line for our imaging purposes. For this line, both the VTA and frontal cortex were examined and compared to the labeling seen in our virally injected CX3CR1^GFP^/Th-Cre mice. Sample images of both our injected CX3CR1^GFP^/Th^Cre^ and CX3CR1^GFP^/DAT-Cre/Ai14 mice were collected as z stacks with a step size of 1µm using the confocal microscope. Due to the profound absence of mesofrontal dopaminergic axon labeling in our CX3CR1^GFP^/DAT-Cre/Ai14mice and the ectopic labeling of cortical neurons, no formal quantification was done using this line.

### Statistics

Statistical analyses were performed in Prism 9 (GraphPad, La Jolla, CA). All n-values represent individual animals and our sample sizes were based on the norms for similar experiments found in the literature^3,8,9^. All individual values are represented on our plots with the mean± s.e.m. included. For all analyses, α=0.05. Statistical differences between two means were calculated using two-tailed unpaired, or paired t-tests. Statistical differences between multiple means were calculated using one-way or two-way ANOVAs with Bonferroni, Holm-Šídák, Šídák or Dunnett’s post hoc comparisons where appropriate. In cohorts with multiple time points, with occasional missing time points, Mixed-effects model (REML) Fixed effects (type III) was used in place of ANOVA to allow for missing values.

