## Supplemental Figures for "Dopaminergic signaling regulates microglial surveillance and adolescent plasticity in the frontal cortex"

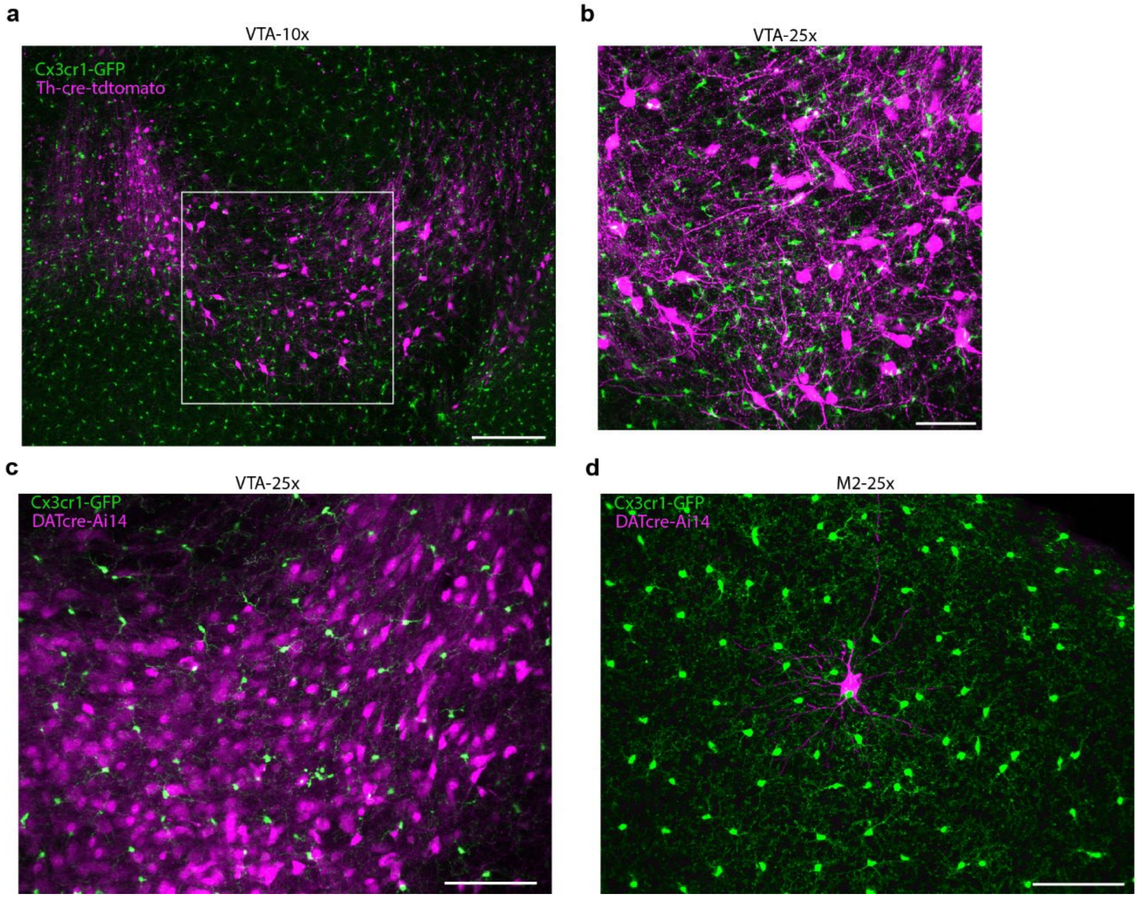


**Supplemental Figure 1| Cx3cr1$^{GFP}$/Th-Cre mice are more effective for imaging the mesofrontal DA circuit than Cx3cr1$^{GFP}$/DAT-Cre/Ai14 mice.** (**a**) Representative maximum intensity Z-projection of a 10x confocal image of VTA in Cx3cr1$^{GFP}$/Th-Cre mice with AAV-CAG-FLEX-tdTomato injection into the VTA (Th+ neurons-magenta, microglia-green) (**b**) Maximum intensity Z-projection of a 25x image of the square ROI from panel a. (**c**) Representative image of the VTA from a Cx3cr1$^{GFP}$/DAT-Cre/Ai14 mouse (DAT+ neurons-magenta, microglia-green). (**d**) Representative image of M2 frontal cortex in a Cx3cr1$^{GFP}$/DAT-Cre/Ai14 mouse. Note the lack of axonal labeling and the presence of an ectopically labelled cortical neuron. (DAT+ neuron-magenta, microglia-green). Scale bars 100μm (a,c,d) and 80μm (b).

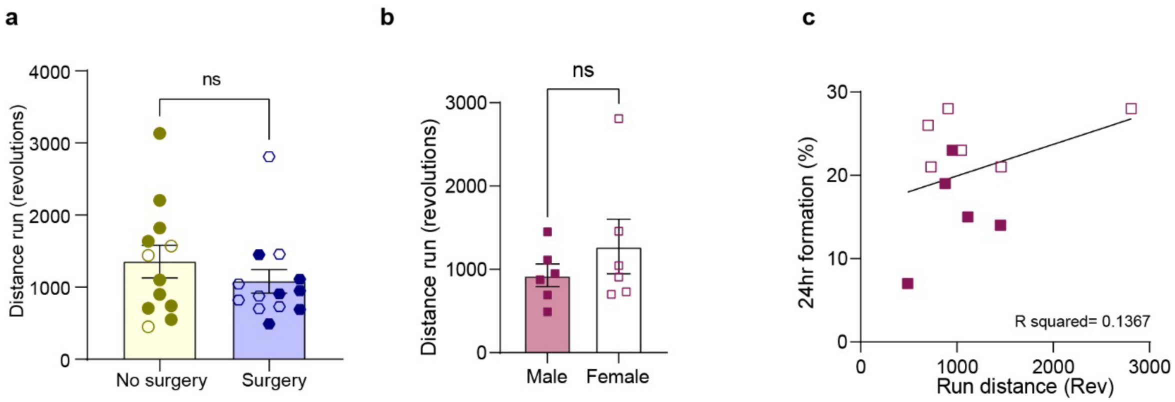


**Supplemental Figure 2| Chronic cranial window preparations do not alter wheel running behavior** (**a**) Chronic cranial window implantation does not significantly alter wheel running (n: No surgery=12, surgery=13, two-tailed unpaired t-test, p=0.3306, t(23)=0.9939). (**b**) Females do not run significantly more than males (n: male=6, female=6; two-tailed unpaired t-test, p=0.3513, t(10)=0.9777). (**c**) There appears to be a positive correlation between run distance and bouton formation at 24hrs. However, the correlation does not reach significance (linear regression slope significance, p=0.2631). One of the males included in b did not have a 24hr imaging time point and thus is excluded in c. Graph shows mean $\pm$ S.E.M. Individual points represent individual animals with females as hollow symbols and males as solid symbols.

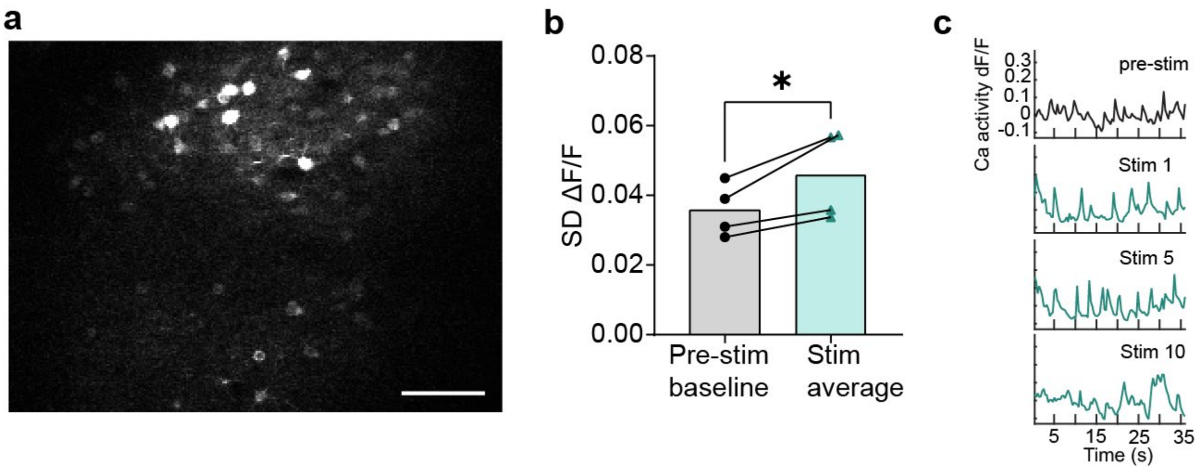


**Supplemental Figure 3| Phasic optogenetic stimulation of the mesofrontal DA axons increases cortical neuronal activity. (**a**)** Representative frame from an *in vivo* two-photon calcium imaging session. M2 frontal cortical neurons were labeled with AAV9-CAMKII-GCaMP6s injected locally. (**b**) Phasic optogenetic stimulation of the mesofrontal DA axons (473nm, 20mW output, 50Hz pulse train, 3ms/pulse, 10 pulses/train, 1 train/min for 10min) elicits a significant change in calcium activity in M2 neurons. Calcium imaging was conducted (0.367s/frame, 100 frames) just prior to stimulation (pre-stim baseline) and in between each of the ten pulses. The standard deviation (SD) of the calcium activity (dF/F) trace in those ten responses were averaged and compared to that in the pre-stim baseline. (n=4 mice, two-tailed paired t-test, p=0.0472, t(3)=3.259). (**c**) Example traces from just prior to stimulation (pre-stim) and between stimulation pulses. Graph shows mean ± S.E.M. Individual points represent individual animals.

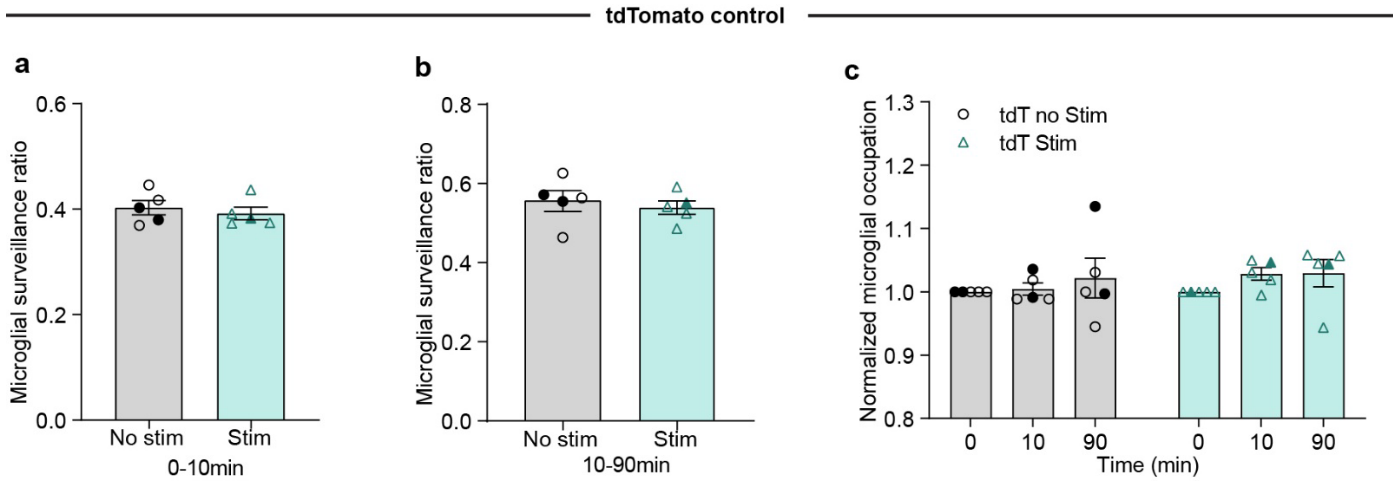


**Supplemental Figure 4| In the absence of channelrhodopsin expression, the phasic light stimulation paradigm does not alter adolescent microglial dynamics.** (**a**) Microglial surveillance is unchanged during phasic light stimulation in the absence of channelrhodopsin (n=5 mice, two-tailed unpaired t-test, p=0.5553, t(8)=0.6155). (**b**) Microglial surveillance is unchanged after phasic light stimulation in the absence of channelrhodopsin (n=5 mice, two-tailed unpaired t-test, p=0.6068, t(8)=0.5355). (**c**) Microglial occupation is not altered by phasic light stimulation in the absence of channelrhodopsin (n=5 mice, two-way repeated measures ANOVA, time x stimulation p=0.7867, F(2,16)=0.2436). Graphs show mean ± S.E.M. No stim and stim experiments were conducted within the same animals. Individual points represent individual animals with females as hollow symbols and males as solid symbols.

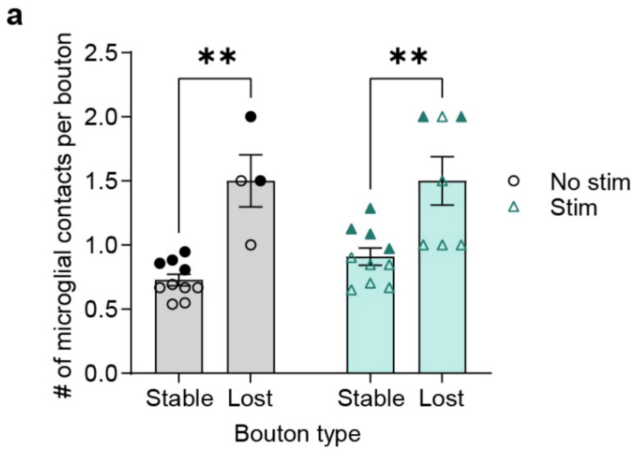


**Supplemental Figure 5| Microglia contact eliminated boutons more frequently than stable boutons.** (**a**) Eliminated boutons are contacted more frequently by microglia than stable boutons (Mixed-effects model, Fixed effects [type III], bouton type p=0.0007, F(1,16)=17.57, Holm-Šídák's multiple comparisons, No stim Stable v. Lost p=0.0010, Stim Stable v. Lost p=0.0012). Graph shows mean ± S.E.M. **p<0.01. No stim and stim experiments were conducted within the same animals. Individual points represent individual animals with females as hollow symbols and males as solid symbols.

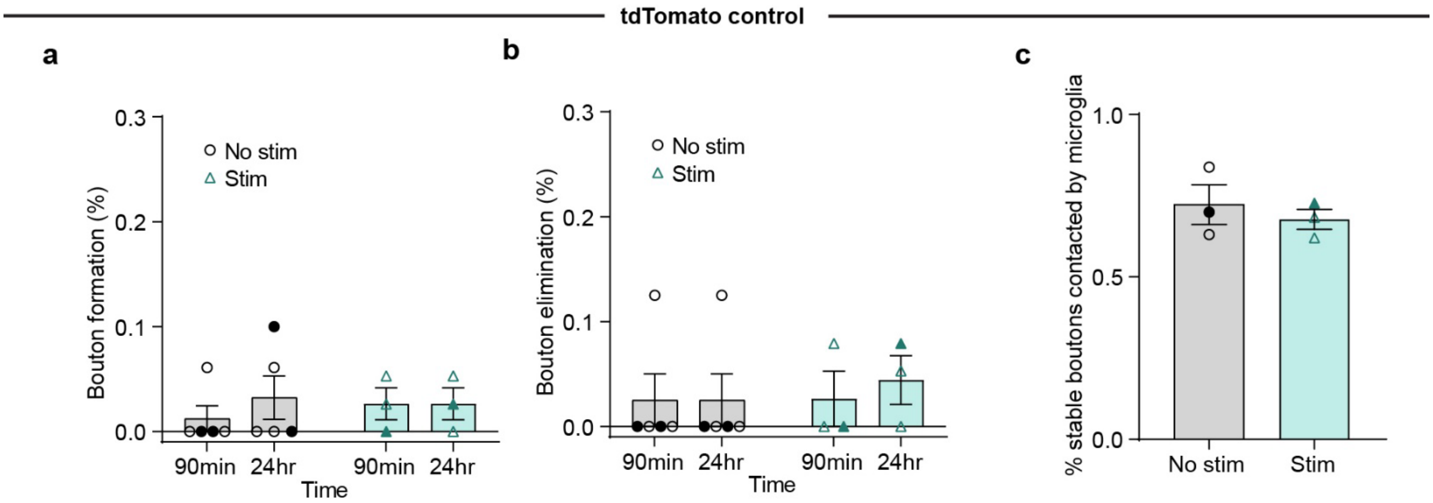


**Supplemental Figure 6| In the absence of channelrhodopsin expression, the phasic light stimulation paradigm does not alter adolescent mesofrontal plasticity or microglial dynamics.** (**a**) Phasic light stimulation does not alter bouton formation rates in the absence of channelrhodopsin (n=5 mice, two-way repeated measures ANOVA, time x stimulation p=0.5661, F(1,6)=0.3684). (**b**) In the absence of channelrhodopsin, phasic light stimulation does not alter bouton elimination rates (n=5 mice, two-way repeated measures ANOVA, time x stimulation P=0.6385, F (1, 6) = 0.2446) (**c**) In the absence of channelrhodopsin, microglial contacts with stable boutons are not altered by phasic light stimulation (n=3 mice, two-tailed unpaired t-test, p=0.5418, t(4)=0.6661) Graphs show mean ± S.E.M. No stim and stim experiments were conducted within the same animals. Individual points represent individual animals with females as hollow symbols and males as solid symbols.

a

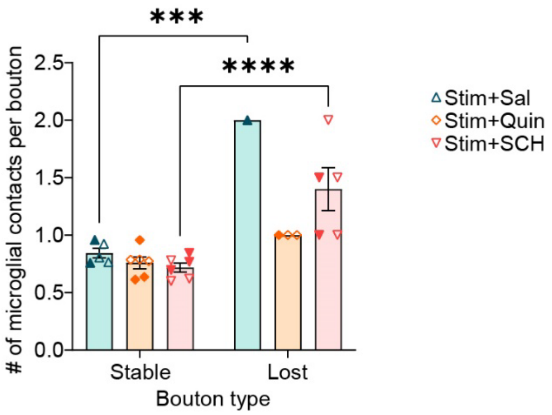


**Supplemental Figure 7| Microglia make more contacts with eliminated boutons than stable boutons in adolescence.** Mixed-effects model, Fixed effects [type III] interaction $p=0.0079$ $F(2,20)=6.232$, Šídák's multiple comparisons Stable v. Lost Stim+Sal $p=0.0002$, Stim+SCH $p<0.0001$. Graph shows mean ± S.E.M, ***$p<0.005$, ****$p<0.0001$. Individual points represent individual animals with females as hollow symbols and males as solid symbols.

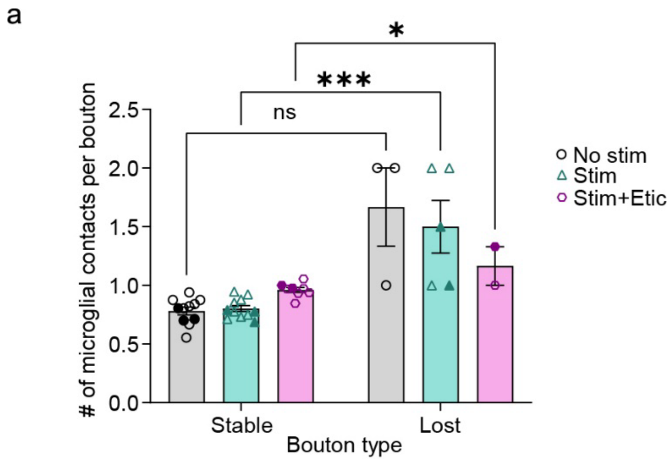


**Supplemental Figure 8| In adults microglia make more frequent contacts with eliminated boutons than stable boutons.** Mixed-effects model, Fixed effects [type III] interaction $p<0.0001$ $F(2,17)=34.21$, Šídák's multiple comparisons Stable v. Lost: Stim $p=0.0004$, Stim+Etic $p<0.0312$. Graph shows mean $\pm$ S.E.M *$p<0.05$, ***$p<0.005$. No stim, Stim and Stim+Etic experiments were conducted within the same animals. Individual points represent individual animals with females as hollow symbols and males as solid symbols.

**Supplemental Video 1**

Time-lapse movie taken through a chronic cranial window showing the surveillance of adolescent microglia in the M2 frontal cortex of awake unstimulated mice taken over 10min at 1min intervals (10µm maximum intensity z-projections were compressed at each time point, video representative of observations made in n=11 mice) Scale bar, 20µm.

**Supplemental Video 2**

Time-lapse movie taken through a chronic cranial window showing the reduced surveillance of adolescent microglia in the M2 frontal cortex of awake mice in response to phasic optogenetic stimulation of DA axons taken over 10min at 1min intervals (10µm maximum intensity z-projections were compressed at each time point, video representative of observations made in n=11 mice) Scale bar, 20µm.

**Supplemental Video 3**

Time-lapse movie taken through a chronic cranial window showing the surveillance of adolescent microglia in the M2 frontal cortex of awake unstimulated mice taken over 80min at 10min intervals (30µm maximum intensity z-projections were compressed at each time point, video representative of observations made in n=11 mice) Scale bar, 20µm.

**Supplemental Video 4**

Time-lapse movie taken through a chronic cranial window showing the increased surveillance of adolescent microglia in the M2 frontal cortex of awake mice after phasic optogenetic stimulation of DA axons taken over 80min at 10min intervals (30µm maximum intensity z-projections were compressed at each time point, video representative of observations made in n=11 mice) Scale bar, 20µm.

**Supplemental Video 5**

Time-lapse movie taken through a chronic cranial window showing putative contacts between microglial processes and the axon backbone in the M2 frontal cortex of awake adolescent mice in response to phasic optogenetic stimulation of DA axons taken over 90min at 10min intervals and including the final 24hr time point (5µm maximum intensity z-projections presented to assist in visualizing contacts though analysis was conducted on individual slices, video representative of observations made in n=10 mice) Box highlights region of interest. Scale bar, 20µm.

**Supplemental Video 6**

Time-lapse movie taken through a chronic cranial window showing the reduced surveillance of adolescent microglia in the M2 frontal cortex of awake control mice dosed with Sal and receiving phasic optogenetic stimulation of DA axons taken over 10min at 1min intervals (10µm maximum intensity z-projections were compressed at each time point, video representative of observations made in n= 5 mice) Scale bar, 20µm.

**Supplemental Video 7**

Time-lapse movie taken through a chronic cranial window showing the reduced surveillance of adolescent microglia in the M2 frontal cortex of awake Quin dosed mice receiving phasic optogenetic stimulation of DA axons taken over 10min at 1min intervals (10µm maximum intensity z-projections were compressed at each time point, video representative of observations made in n= 6 mice) Scale bar, 20µm.

**Supplemental Video 8**

Time-lapse movie taken through a chronic cranial window showing that SCH inhibits adolescent microglial retraction in response to phasic optogenetic stimulation of DA axons taken over 10min at 1min intervals (10µm maximum intensity z-projections were compressed at each time point, video representative of observations made in n= 6 mice) Scale bar, 20µm.

**Supplemental Video 9**

Time-lapse movie taken through a chronic cranial window showing the increased surveillance of adolescent Sal dosed microglia in the M2 frontal cortex of awake mice after phasic optogenetic stimulation of DA axons taken over 80min at 10min intervals (30µm maximum intensity z-projections were compressed at each time point, video representative of observations made in n=5 mice) Scale bar, 20µm.

**Supplemental Video 10**

Time-lapse movie taken through a chronic cranial window showing that Quin prevents the increased surveillance of adolescent microglia in the M2 frontal cortex of awake mice after phasic optogenetic stimulation of DA axons taken over 80min at 10min intervals (30µm maximum intensity z-projections were compressed at each time point, video representative of observations made in n=6 mice) Scale bar, 20µm.

**Supplemental Video 11**

Time-lapse movie taken through a chronic cranial window showing that SCH dosing maintains microglia in a high state of surveillance in the M2 frontal cortex of awake adolescent mice after phasic optogenetic stimulation of DA axons taken over 80min at 10min intervals (30µm maximum intensity z-projections were compressed at each time point, video representative of observations made in n=6 mice) Scale bar, 20µm.

**Supplemental Video 12**

Time-lapse movie taken through a chronic cranial window showing the surveillance of adult microglia in the M2 frontal cortex of awake unstimulated mice taken over 10min at 1min intervals (10µm maximum intensity z-projections were compressed at each time point, video representative of observations made in n=11 mice) Scale bar, 20µm.

**Supplemental Video 13**

Time-lapse movie taken through a chronic cranial window showing the reduced surveillance of adult microglia in the M2 frontal cortex of awake mice in response to phasic optogenetic stimulation of DA axons taken over 10min at 1min intervals (10µm maximum intensity z-projections were compressed at each time point, video representative of observations made in n= 11 mice) Scale bar, 20µm.

**Supplemental Video 14**

Time-lapse movie taken through a chronic cranial window showing the reduced surveillance of adult microglia in the M2 frontal cortex of awake Etic dosed mice in response to phasic optogenetic stimulation of DA axons taken over 10min at 1min intervals (10µm maximum intensity z-projections were compressed at each time point, video representative of observations made in n= 11 mice) Scale bar, 20µm.

**Supplemental Video 15**

Time-lapse movie taken through a chronic cranial window showing the surveillance of adult microglia in the M2 frontal cortex of awake unstimulated mice taken over 80min at 10min intervals (30µm maximum intensity z-projections were compressed at each time point, video representative of observations made in n=11 mice) Scale bar, 20µm.

**Supplemental Video 16**

Time-lapse movie taken through a chronic cranial window showing the recovery of surveillance post-stimulation in adult microglia in the M2 frontal cortex of awake mice after phasic optogenetic stimulation of DA axons taken over 80min at 10min intervals (30µm maximum intensity z-projections were compressed at each time point, video representative of observations made in n=11 mice) Scale bar, 20µm.

**Supplemental Video 17**

Time-lapse movie taken through a chronic cranial window showing the increased surveillance of adult Etic dosed microglia in the M2 frontal cortex of awake mice after phasic optogenetic stimulation of DA axons taken over

80min at 10min intervals (30μm maximum intensity z-projections were compressed at each time point, video representative of observations made in n=9 mice) Scale bar, 20μm.
